# Histone H3 clipping is a novel signature of human neutrophil extracellular traps

**DOI:** 10.1101/2021.03.15.434949

**Authors:** Dorothea O Tilley, Ulrike Abuabed, Ursula Zimny Arndt, Monika Schmid, Stefan Florian, Peter R. Jungblut, Volker Brinkmann, Alf Herzig, Arturo Zychlinsky

## Abstract

Neutrophils are critical to host defence, executing diverse strategies to perform their antimicrobial and regulatory functions. One tactic is the production of neutrophil extracellular traps (NETs). In response to certain stimuli neutrophils decondense their lobulated nucleus and release chromatin into the extracellular space through a process called NETosis. However, NETosis, and the subsequent degradation of NETs, can become dysregulated. NETs are proposed to play a role in infectious as well as many non-infection related diseases including cancer, thrombosis, autoimmunity and neurological disease. Consequently, there is a need to develop specific tools for the study of these structures in disease contexts. In this study, we identified a NET-specific histone H3 cleavage event and harnessed this to develop a cleavage site-specific antibody for the detection of human NETs. By microscopy, this antibody distinguishes NETs from chromatin in purified and mixed cell samples. It also detects NETs in tissue sections. We propose this antibody as a new tool to detect and quantify NETs.

## Introduction

Neutrophil extracellular traps (NETs) are extracellular structures consisting of chromatin components, including DNA and histones, and neutrophil proteins (Brinkmann et al., 2004, Urban et al., 2009). NETs were first described as an antimicrobial response to infection, facilitating trapping and killing of microbes (Brinkmann et al., 2004). They are found in diverse human tissues and secretions where inflammation is evident (recently reviewed by Sollberger et al., 2018). NETs are produced in response to a wide-range of stimuli; bacteria (Brinkmann et al., 2004); fungi (Urban et al., 2006); viruses (Saitoh et al., 2012; Schonrich et al., 2015); crystals (Schauer et al., 2014); and mitogens (Amulic et al., 2017). Both NADPH oxidase (NOX) dependent and NOX independent mechanisms lead to NET formation (Bianchi et al., 2009; Hakkim et al., 2011; Kenny et al., 2017; Neeli and Radic, 2013). NETs are also observed in sterile disease, including multiple types of thrombotic disease (recently reviewed by Jimenez-Alcazar et al., 2017) and even neurological disease (Zenaro et al., 2015). NETs, or their components, are implicated in the development and exacerbation of autoimmune diseases including psoriasis, vasculitis and systemic lupus erythematosus (recently reviewed by Papayannopoulos, 2018) as well as cancer and cancer metastasis (Albrengues et al., 2018; Cools-Lartigue et al., 2013; Demers et al., 2016). Consequently, there is an urgency across multiple fields to establish the pathological contribution of NETs to disease. However, the detection of NETs in affected tissues remains a challenge.

NETs are histologically defined as areas of decondensed DNA and histones that colocalise with neutrophil granular or cytoplasmic proteins. Thus, reliable detection of NETs requires a combination of anti-neutrophil and anti-chromatin antibodies as well as DNA stains. Immunofluorescent microscopy is a useful method to detect NETs in tissue sections and *in vitro* experiments. However, this can be challenging since NET components are distributed across the large decondensed structure resulting in a weak signal. For example, the signal of antibodies to neutrophil elastase (NE) is significantly dimmer in NETs than in the granules of resting cells where this protease is highly concentrated. Conversely, anti-histone antibodies stain NETs strongly but not nuclei of naïve neutrophils, where the chromatin is compact and less accessible. This differential histone staining property can be exploited for the detection and quantification of NETs (Brinkmann et al., 2012). However, sample preparation and the subsequent image analysis make results between different labs difficult to compare. Thus, there is a need to identify antibodies against NET antigens.

NETs can also be detected through post-translational modifications (PTMs) that occur during NETosis. Histone 3 (H3) is deaminated in arginine residues - the conversion to citrulline (citrullination) - by protein arginine deiminase 4 (PAD4) (Wang et al., 2009). Citrullinated H3 (H3cit) is widely used as a surrogate marker of NETs in both *in vitro* and *in vivo* experiments (Gavillet et al., 2015; Pertiwi et al., 2018; Wang et al., 2009; Yoo et al., 2014; Yoshida et al., 2013). Cleavage of histones by granular derived neutrophil serine proteases (NSPs) also contributes to NETosis (Papayannopoulos et al., 2010). Histone cleavage, or clipping, by cysteine or serine proteases is a bona fide histone PTM that facilitates the gross removal of multiple, subtler, PTMs in the histone tail and is conserved from yeast to mammals (Dhaenens et al., 2015). Until now, histone clipping has not been exploited for the detection of NETs but recent work by our group showed that histone H3 cleavage is a conserved response to diverse NET stimuli, including *Candida albicans* and Group B Streptococcus (Kenny 2017). Thus, in this study we map the site(s) of histone H3 cleavage during NETosis. We developed a new monoclonal antibody against cleaved H3 that detects human NETs *in vitro* and in histological samples. This antibody also facilitates easier NET quantification.

## Results

### Serine protease dependent cleavage of Histone H3 N-terminal tails during NET formation

Histones are processed in response to phorbol 12 myristate 13 acetate - PMA (Papayannopoulos et al., 2010; Urban et al., 2009) and other NET stimuli (Kenny et al., 2017). Indeed, the cleavage products of H3 were consistent between stimuli (Kenny et al., 2017). This suggests that H3 proteolysis occurs at specific sites during NETosis. In a time course experiment of human neutrophils incubated with PMA, we detected a H3 cleavage product of ∼14 kDa as early as 30 min post-stimulation (Figure 1A). Further cleavages occurred between 60 and 90 min, yielding products of approximately 13 kDa and 10 kDa, respectively. The histone N-terminal tails protrude from the nucleosome core and are a major PTM target (Bannister and Kouzarides, 2011). A C-terminal, but not an N-terminal, histone antibody detected the cleavage products of H3 (Figure 1A). These results indicate that the N-terminus is cleaved in truncated H3.

**Figure 1.**
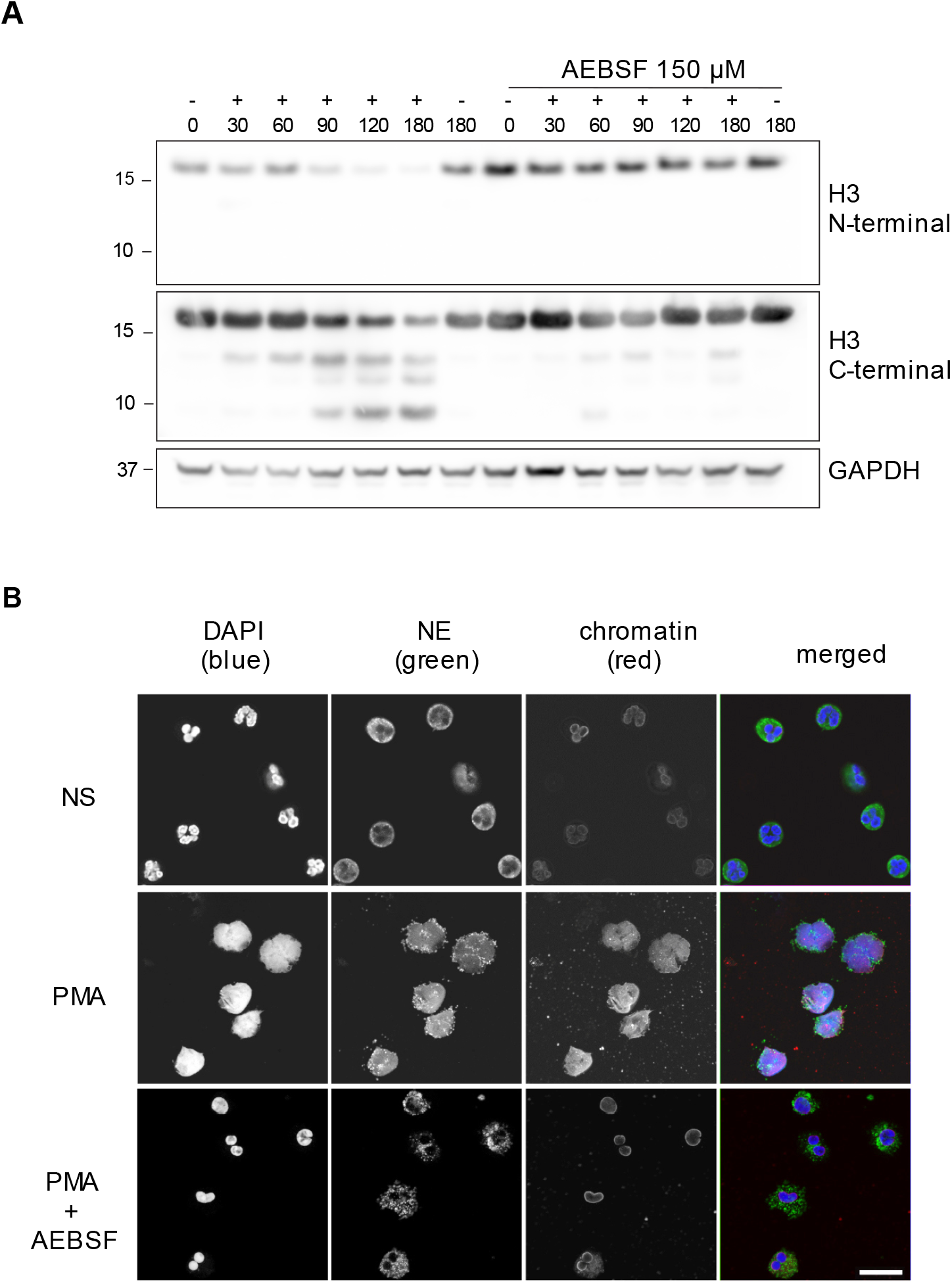
PMA induced Histone H3 cleavage occurs in the N-terminal domain & is prevented by serine protease inhibition. (**A**) Neutrophils were preincubated with the serine protease inhibitor AEBSF for 30 min in microcentrifuge tubes and then stimulated with PMA as indicated in the figure. Lysates were resolved by SDS-PAGE and immunoblotted with N- and C-terminal antibodies to H3. GAPDH was used as a loading control. Blots representative of 3 independent experiments. (**B**) Immunofluorescent confocal microscopy of NET formation. Neutrophils were seeded on coverslips and preincubated with AEBSF before stimulation with PMA (50 nM) as indicated in the figure. At 150 min the cells were fixed and stained for neutrophil elastase (NE), chromatin (using a H2A-H2B-DNA antibody PL2.3) and DNA (DAPI [4′,6-diamidino-2-phenylindole]). NS: non-stimulated. Images were taken at 63X and the scale bar is 20 µm. Images are representative of 3 independent experiments.

Neutrophil azurophilic granules are rich in serine proteases that can degrade histones *in vitro* (Papayannopoulos et al., 2010). We tested the contribution of these proteases to H3 cleavage during NET formation. Preincubation with the serine protease inhibitor, AEBSF (4- [2-aminoethyl] benzensulfonylfluoride) (Figure 1A), but not with the cysteine protease inhibitor E64 (Figure 1– figure supplement 1), inhibited H3 cleavage upon PMA stimulation. AEBSF also inhibited NET formation and nuclear decondensation as shown by immunofluorescent microscopy (Figure 1B). PMA induced NETosis requires NADPH oxidase activity and, at high concentrations, AEBSF can inhibit activation of NAPDH oxidase (Diatchuk et al., 1997). To rule out this upstream effect, we showed that AEBSF did not inhibit ROS production at the concentrations used in our assay (Figure 1–figure supplement 2). Similarly, we verified that at these concentrations AEBSF was not cytotoxic, as shown by limited LDH release (Figure 1–figure supplement 3). This data shows that serine proteases cleave the N-terminus of H3 early during NET formation.

### Histone H3 is cleaved at a novel site in the globular domain

To identify the precise H3 cleavage sites we prepared histone enriched extracts from primary neutrophils stimulated with PMA for 90 min and then purified H3 by RP-HPLC as previously described (Shechter et al., 2007). A schematic summary of this is presented in Figure 2-figure supplement 1. We identified the fractions containing H3 and its cleaved products by Western blot with anti-H3 C-terminal antibodies (Figure 2A). As expected, H3 was the last core histone to elute (at 45-46 min).

**Figure 2.**
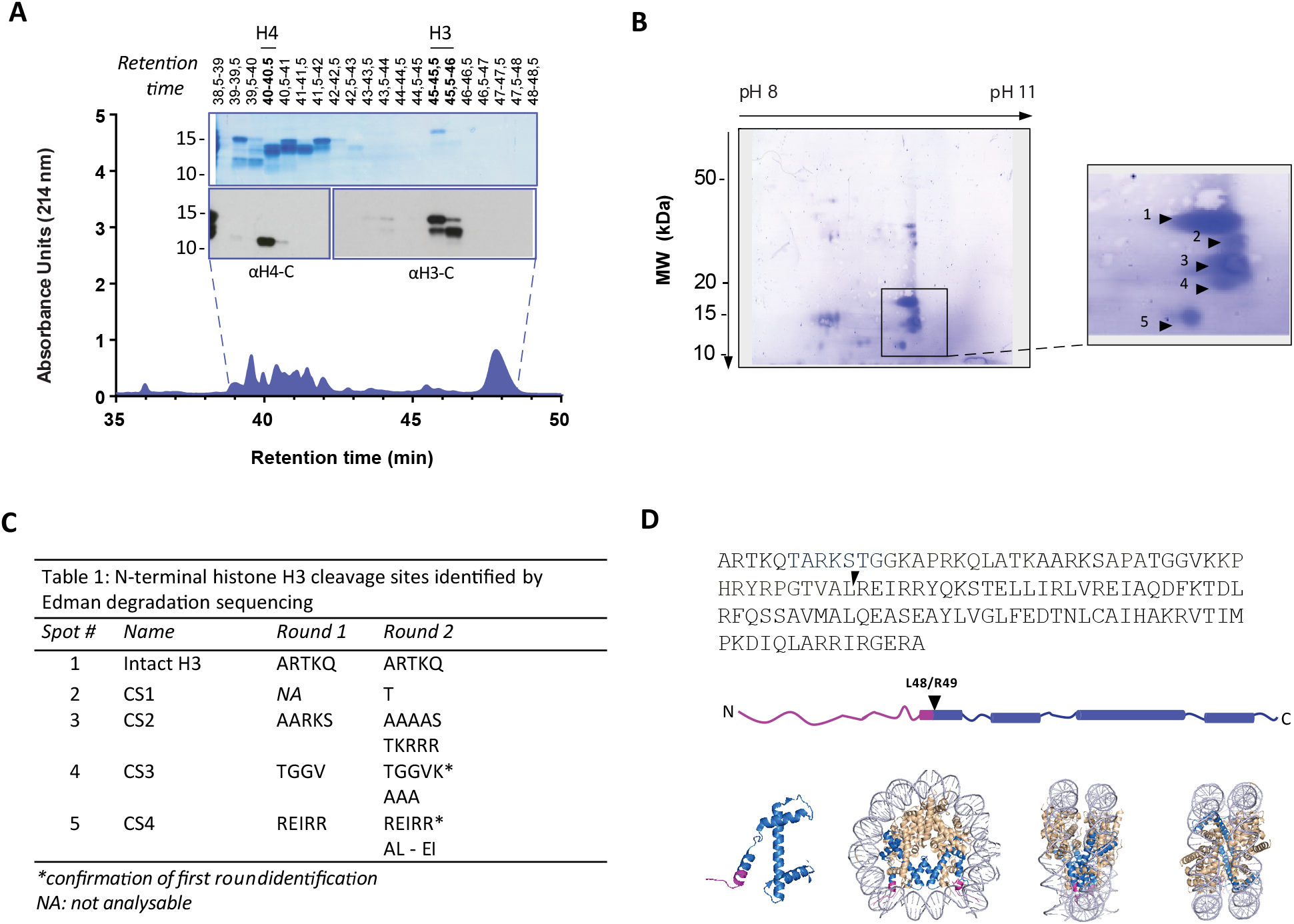
Identification of histone H3 cleavage sites in NET formation. (**A**) Representative RP-HPLC chromatogram of acid extracted histones from NETs and corresponding 1D-SDS-PAGE and immunoblots to identify H3 and H4 containing fractions. Histone enriched supernatants were prepared from neutrophils stimulated with PMA for 90 min. Purification and subsequent 2-DE analysis was repeated 3 times with independent donors. (**B**) Representative Coomassie stained blot of pooled H3 fractions separated by 2-DE. Inset is a zoom of all spots (1-5) identified as histone H3 by mass spectrometry. Other proteins identified are listed in Figure 2-figure supplements 2 and 3 (**C**) Summary of Edman degradation sequencing results of the H3 spots in two independent experiments. In the second experiment overlapping sequences were detected. However, the detected amino acids for spot 4 and 5 confirmed the initial sequence identification from round 1*. (**D**) Schematic representation of the cleavage site of the truncated H3 product in both the linear sequence of H3 and in the nucleosomal context. H3 is represented in blue and the pink tail region and partial alpha helix represent the part of H3 that is removed. The nucleosome structure is adapted from PDB 2F8N (Chakravarthy, 2005).

We further separated H3 and its truncated forms by two-dimensional electrophoresis (2-DE) and confirmed their identity by mass spectrometry (Figure 2B and Figure 2-figure supplements 2 and 3). The sequence coverage did not include residues that allowed the differentiation of H3 variants. The N-terminals of the separated H3 fragments were not covered by MS and therefore sequenced by Edman degradation from 2 independent experiments (Figure 2C). The N-terminal sequence of the largest molecular weight H3 spot matched that of intact H3 (Figure 2B, spot 1). We did not obtain reliable sequencing of spots 2 and 3 but the cleavage sites of spots 4 and 5 were identified. The most truncated H3 fragment (spot 5) was cleaved between L48 and R49, in the globular domain of the protein, within the nucleosome core structure (Figure 2D). This is a previously unidentified cleavage site in H3 and, thus, we selected cleavage at H3R49 as a candidate marker of NETs.

### Generation of a histone H3 cleavage site monoclonal antibody

We adopted a similar strategy to Duncan and colleagues (2008) to raise antibodies against the cleaved site. We designed a lysine branched immunogen containing the 5 amino acids at the carboxylic side of the H3R49 cleavage site (outlined in Figure 3-figure supplement 1 and 2) and used it to immunize mice. After preliminary screening by ELISA against the immunogen and control peptides, we selected sera, and later hybridoma clones, that detected cleaved H3 in immunoblots of PMA stimulated cell lysates. We excluded sera and clones that detected full length histone H3 in addition to cleaved H3 (Figure 3A). We selected sera and clones that detected NETs but not resting chromatin of naive neutrophils by immunofluorescence microscopy. Of the 6 mice immunised, we obtained one stable clone, 3D9, that functioned in both Western blot and microscopy – other clones performed only in Western blot (data not shown). 3D9 recognised a protein of ∼10 kDa in neutrophils stimulated with PMA for 120 min and longer but did not detect any protein in resting or early stimulated cells (Figure 3A). This band corresponded in size with the smallest H3 fragment detected by the H3 C-terminal antibody. Interestingly, by microscopy, 3D9 exclusively recognised neutrophils undergoing NETosis – with decondensing chromatin (Figure 3B). To validate the specificity of the antibody for the *de novo* N-terminal H3 epitope of NETs, we performed competition experiments with the immunising peptide and demonstrated that it could block 3D9 binding to NETs as shown by immunofluorescent microscopy (Figure 3-figure supplement 3).

**Figure 3.**
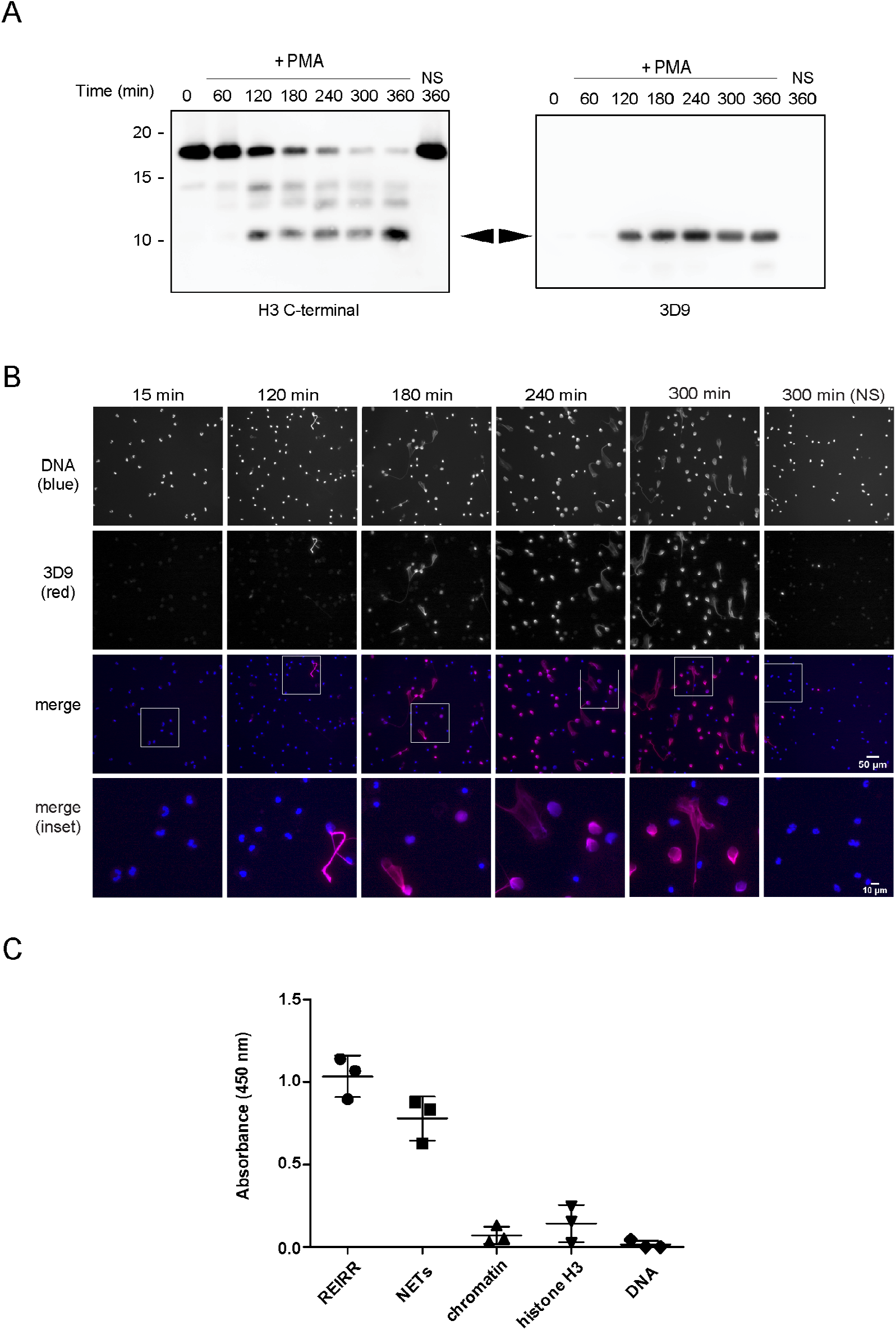
Screening and detection of cleaved H3 & NETs by 3D9. (**A**) Immunoblots of lysates prepared from neutrophils stimulated with PMA (50 nM) for the times indicated in the figure. H3 C-terminal antibody was used as a control to detect all H3 forms while a single band (cleaved H3) was detected by the newly generated monoclonal antibody, 3D9. (**B**) Immunofluorescent microscopy of neutrophils stimulated with PMA and fixed at the indicated times. Samples were stained with Hoechst (DNA - blue) and 3D9 (with Alexafluor-568 conjugated secondary antibody - red). NS: non-stimulated. Images were taken on an upright fluorescent microscope at 20X. Scale bars – 50 µm (full field) and 10 µm (inset) (**C**) Direct ELISA for cleaved H3 in NETs, chromatin (A549 lung epithelial cells), recombinant histone H3 and DNA. Samples were serially diluted and immobilized on a high affinity ELISA plate according to DNA content (for NETs, chromatin and DNA) or protein content (for recombinant histone H3) as determined by PicoGreen and bicinchoninic acid assays respectively. Starting concentration was 1 µg/ml DNA or protein. Cleaved H3 was detected using 3D9 (2 µg/ml) and HRP conjugated anti-mouse secondary antibody and reactions were developed using TMB (3,3’, 5,5’ -tetramethylbenzidine) as a substrate. Data is presented for dilution 200 ng/ml. REIRR peptide control was coated at 20 ng/ml. Data represents mean ± SD of 3 experiments using independent NET donors. Source data can be found in Figure 3-Source data 1.

3D9 binds specifically to cleaved H3. This antibody binds to the immunizing peptide and to isolated NETs, but not to equal concentrations of chromatin, recombinant H3 or purified calf thymus DNA, by direct ELISA (Figure 3C). Furthermore, in immunoprecipitation experiments, 3D9, but not an isotype control, pulled down intact H3 in lysates of naïve and activated neutrophils, but only the cleaved fragment from activated cells (Figure 3-figure supplement 4). We detected these pull downs both by Coomassie and silver stained gels. In a similar experiment, we immunoblotted the immunoprecipitate with the C-terminal antibody as well as 3D9 (Figure 3-figure supplement 4 [ii]), and showed that only our monoclonal antibody recognizes cleaved H3. Together this data shows that 3D9 is selective for cleaved H3 under the denaturing conditions of SDS-PAGE but also recognizes full length H3 under the more native conditions of IP.

### Epitope mapping

To determine the binding site of 3D9 in histone H3, we tested the antibody by ELISA with overlapping linear peptide arrays and helical peptide mimic arrays (listed in Supplementary file 1) based on a sequence (residues 30-70; PATGGVKKPHRYRPGTVALREIRRYQKSTELLIRKLPFQRL) around the H3R49 cleavage site (Figure 4-figure supplement 1 and 2). In such assays acetylation is often used to neutralise the contribution of the amino terminal charge. However, to mimic any potential charge created at the newly revealed N-terminus, R49, we also included arrays of unmodified peptides. Based on overlapping peptides, the putative core epitope in the linear array was (R)EIRR. The peptides ending in REIRR were in all cases in the top 2 of each peptide mimic (Figure 4-figure supplement 3). Interestingly, peptides extended at the N-terminus were still recognized. Moreover, acetylation at the N-terminus of the peptide ending in the REIRR sequence did not affect the binding, suggesting that a free N-terminus may not be recognized by the antibody. We further refined the epitope mapping by amino acid replacement analysis of linear peptides and helical peptide mimetics ending in REIRR (Figure 4-figure supplement 4]). Mutations in Glu51, Ile52, and Arg54 negatively impacted the signal, indicating these residues are critical for epitope recognition. A schematic of the antibody epitope mapped onto H3 is presented in Figure 4.

**Figure 4.**
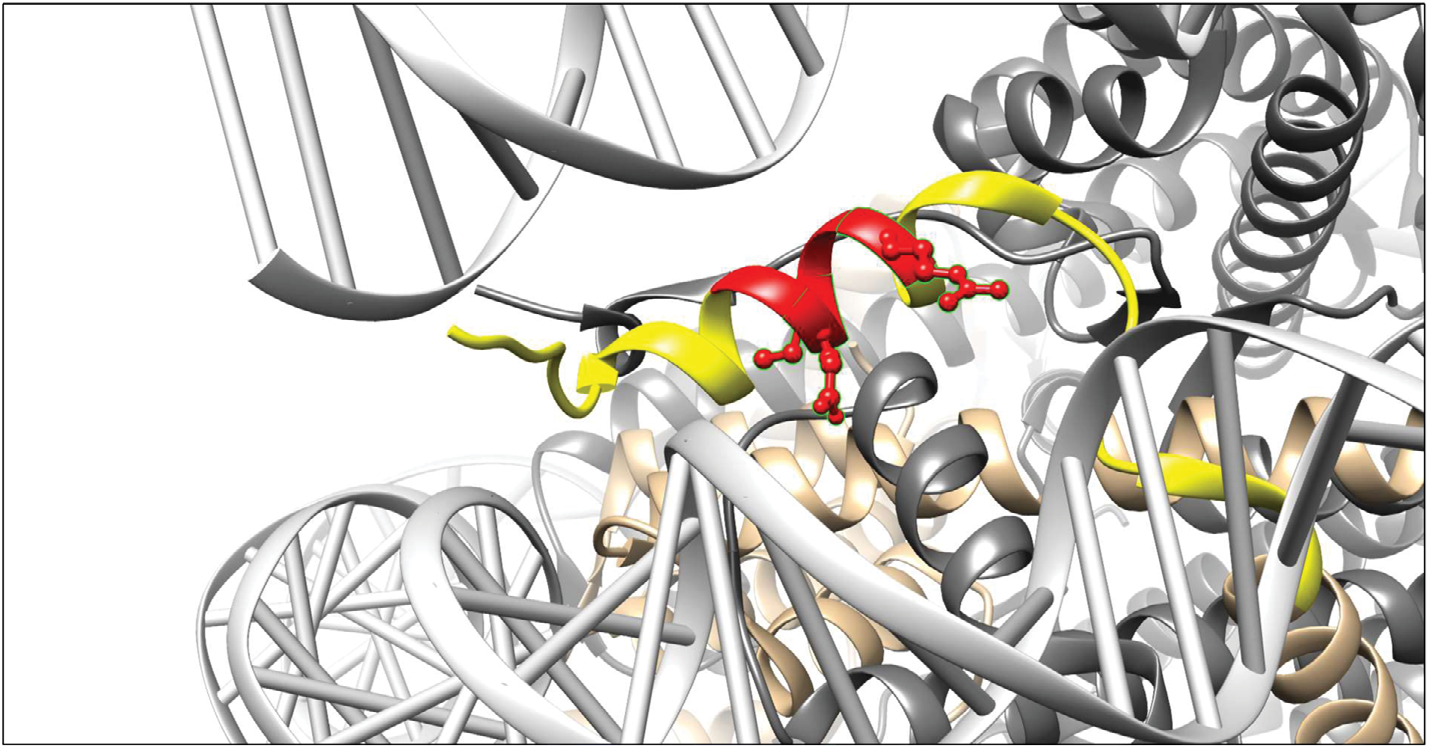
Visualisation of the 3D9 epitope in the nucleosome core complex. Visualization of the putative core epitope for 3D9 mapped on to histone H3 ribbon structure. The observed core binding site of 3D9 to the peptide arrays was depicted on histone H3 (light brown) in the nucleosome complex structure (file 3AZG.pdb). Part of the peptide sequences used in the peptide arrays is coloured in yellow. The core epitope (R)EIRR is displayed in red, with the atoms of the critical residues (Glu51, Ile52, and Arg54) shown. Binding profiles of antibody to linear and helical arrays, in addition to amino acid replacement analysis are presented in Figure 4-figure supplement 1-4.

### Automatic quantification of in vitro generated NETs by microscopy

We tested how 3D9 stained NETs in immunofluorescence in samples that were robustly permeabilized (Triton X-100, 0.5% for 10 min) to facilitate the distribution of the antibody throughout the sample. Figure 5A shows that 3D9 detects decondensed chromatin almost exclusively. In contrast, the anti-chromatin antibody (PL2.3) - directed against a H2A-H2B-DNA epitope (Losman et al., 1992) – stains NETs in addition to condensed nuclei. We compared the staining characteristics of 3D9 and PL2.3 during NET formation. We determined the nuclear area and signal intensity at the indicated time points, from multiple fields of view (Figure 5B). Both antibodies detect the increase in nuclear area characteristic of NETosis between 15 and 180 min after simulation. At later time points, the intensity of PL2.3 staining decreased and failed to discriminate between resting cell nuclei and NETs. In contrast, 3D9 stained nuclei undergoing NETosis with greater intensity than nuclei of non-activated cells.

**Figure 5.**
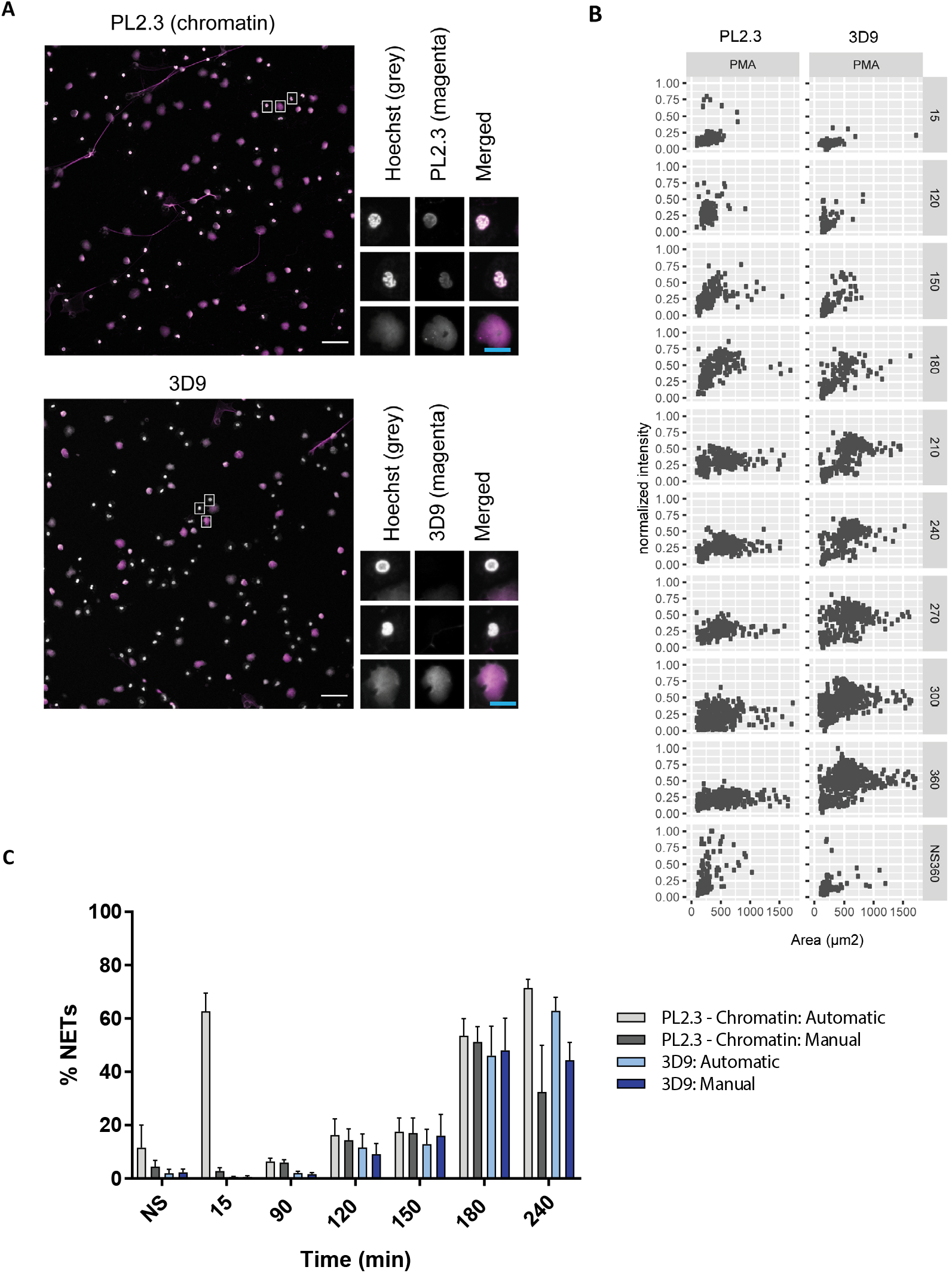
Comparison of NET quantification using an anti-chromatin antibody versus 3D9. Confocal immunofluorescent microscopy of neutrophils stimulated with PMA (180 min) and stained with Hoechst and anti-chromatin antibody (PL2.3) or 3D9. Insets represent selected cells examined at higher magnification (63X) and presented as split channels in grayscale or merged as per the total field of view (20X magnification). White scale bar - 50 µm, cyan scale bar - 10 µm. Images are representative of 3 experiments Comparison of the fluorescent distribution of PL2.3 versus 3D9 staining of PMA stimulated cells over time (6h). Staining intensities were normalized over all images of the respective time course. NS:360: non-stimulated at 360 min. Analysis is performed on one data set that is representative of 3-4 independent time course experiments. (**C**) Comparison of NET quantification using manual or automatic thresholding and segmentation procedures for chromatin antibody (PL2.3) and cleaved H3 antibody (or 3D9). Manual thresholding excludes cells/NETs with a weak signal whereas automatic thresholding includes all objects irrespective of signal. Images for analysis were taken using a fluorescence microscope. Graph represents the mean ± standard deviation, where n=3-5. Source data can be found in Figure 5-Source Data 1.

Publicly available software (ImageJ) can be used to quantify *in vitro* NETosis. We compared 3D9 versus the anti-chromatin antibody with our previously published semi-automatic image analysis (Brinkmann et al., 2012) and with a modified automatic method (Figure 5-figure supplement 1). Both methods use automatic thresholding of the DNA channel (Hoechst) to count total cells/objects. The historical semi-automatic method exploits the differential staining by chromatin antibodies of decondensed chromatin (high signal) over compact chromatin (weak signal) to count cells in NETosis. This method uses a manual thresholding and segmentation procedure (denoted manual in the figure). This manual thresholding step is subject to observer bias. In contrast, the modified method uses automatic thresholding at both stages; total cell and NET counts. Both methods use a size exclusion particle analysis step so that only structures larger than a resting nucleus are counted. Both 3D9 and PL2.3 antibodies effectively quantified NETs using previously published method (Figure 5C - manual). However, PL2.3 failed to accurately quantify the number of NETs with the fully automatic method, specifically at early time points after stimulation (15 min). In this case, the weakly staining lobulated nucleus can extend over a larger surface area during cell activation and adhesion resulting in these cells being wrongly categorised as NETs by the algorithm. Together, this data suggests that the automatic method using 3D9 staining may reduce experimental bias.

### 3D9 detects NETs induced by multiple stimuli

Histone H3 cleavage is a feature of the neutrophil response to multiple NET stimuli (Kenny et al., 2017). 3D9 detects NETs induced by the bacterial toxin nigericin, which induces NETs independently of NADPH oxidase activation (Kenny et al., 2017), by heme in TNF primed neutrophils (Knackstedt et al., 2019) and by the fungal pathogen *Candida albicans* (Figure 6). Interestingly, in *C. albicans* infections, we observed both 3D9 positive and negative NETs.

**Figure 6.**
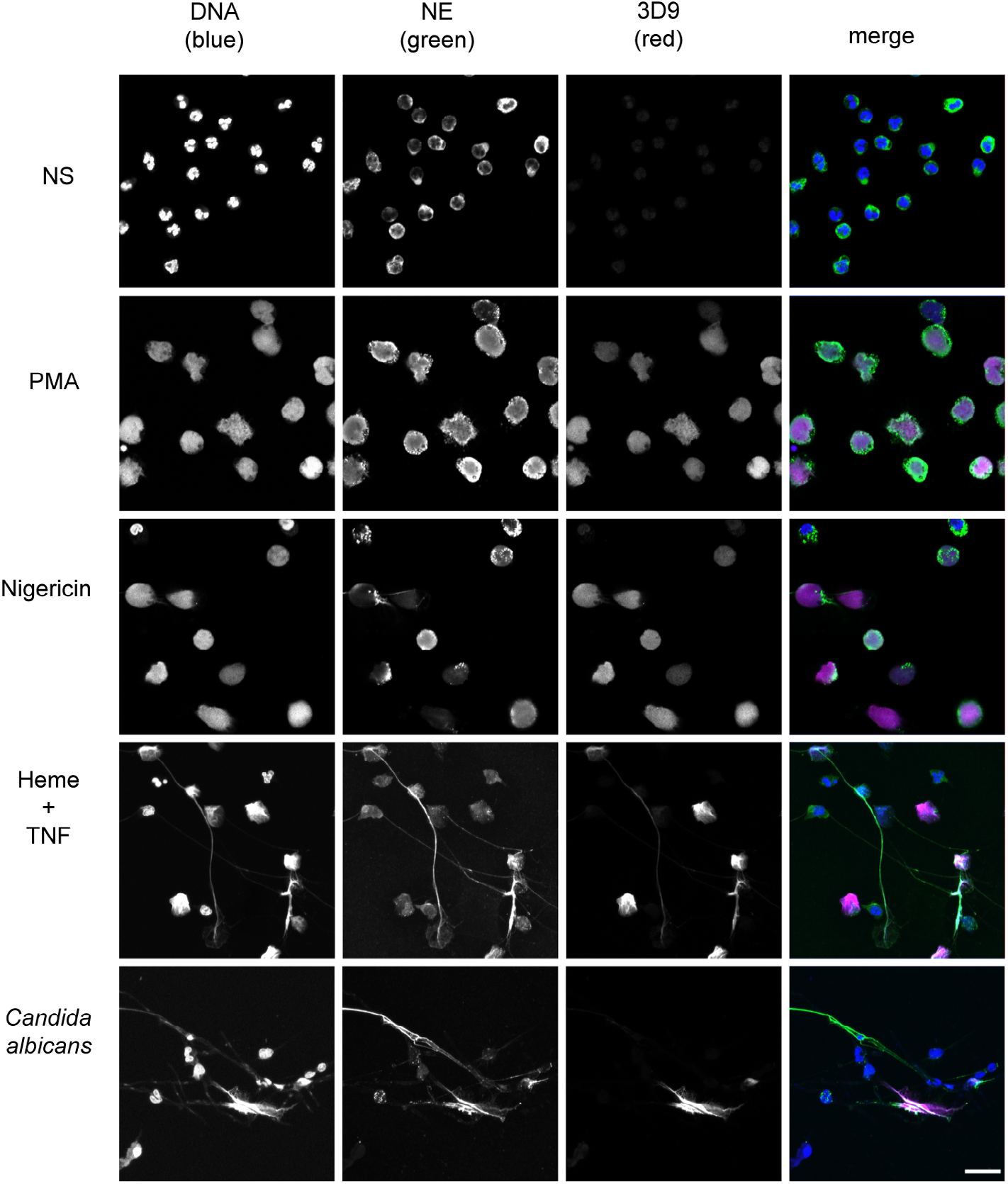
Detection by 3D9 of NETs from diverse stimuli. Immunofluorescence microscopy of neutrophils left unstimulated (NS), stimulated with PMA (50 nM, 2.5h), nigericin (15 µM, 2.5 h), TNF primed and then stimulated with heme (20 µM, 6h), and neutrophils co-cultured with *Candida albicans* hyphae (MOI 5) for 4h. Samples were stained with Hoechst, anti-neutrophil elastase (NE) and 3D9. Scale bar – 50 µm. Images were taken on a confocal microscope at 20X and are representative of 3 experiments with independent donors. Scale bar - 20 µm.

### 3D9 distinguishes NETs in mixed cell samples

Histone H3 clipping, albeit at other sites in the N-terminal tail, was observed in mast cells (Melo et al., 2014) and unstimulated PBMC fractions (Howe and Gamble, 2015). Of note, PBMC fractions often contain contaminating neutrophils (Hacbarth and Kajdacsy-Balla, 1986). To test if 3D9 specifically stained neutrophils treated with NET stimuli, we incubated PBMCs with PMA or nigericin (Figure 7). Importantly, 3D9 detected only nuclei that appeared decondensed in cells that were positive for NE, a specific neutrophil marker. In contrast, the chromatin antibody stained both neutrophils in NETosis and nuclei of other cells. This shows that 3D9 detects NETs specifically even in the presence of other blood cells.

**Figure 7.**
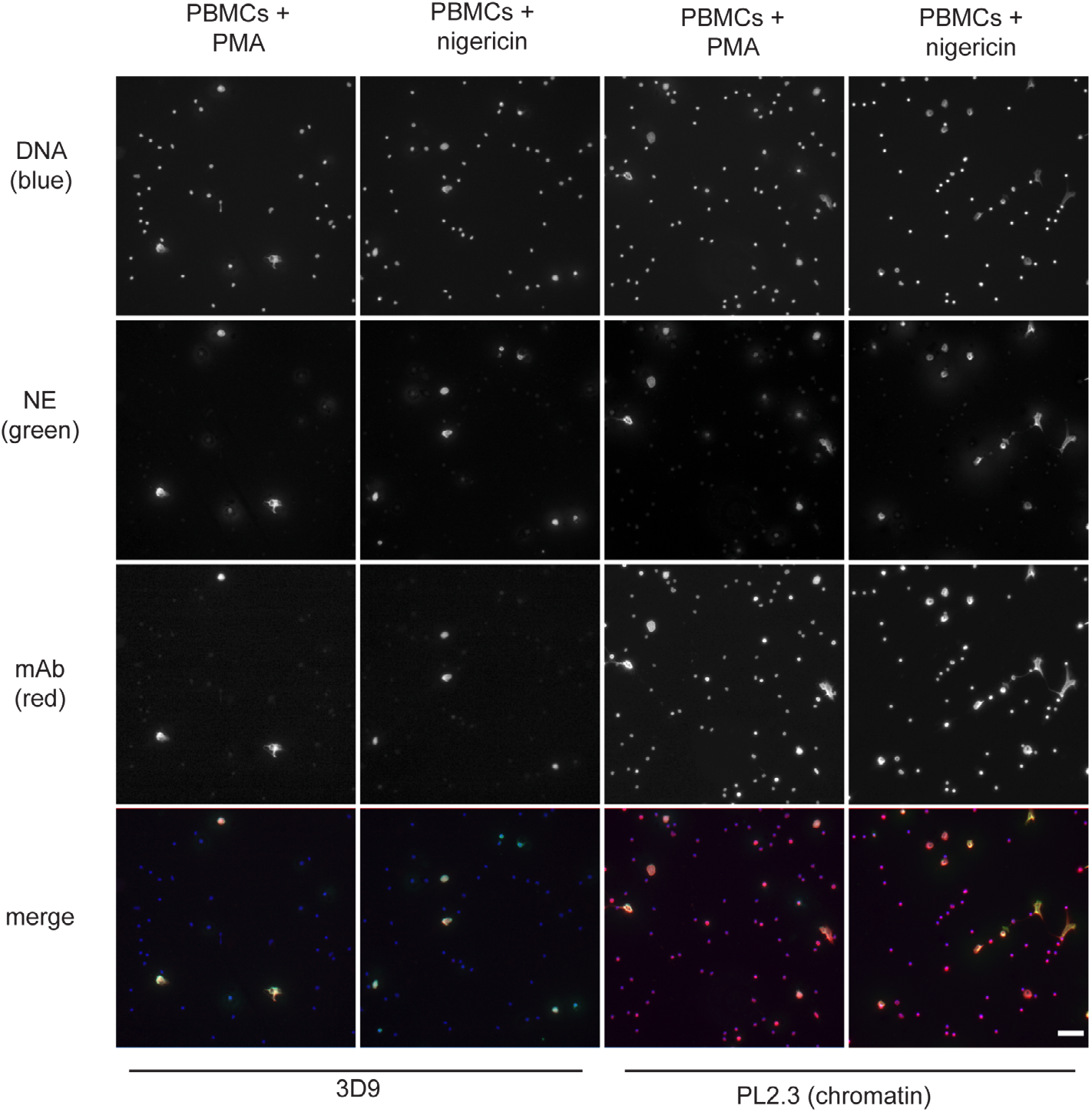
Detection of NETs in mixed cell fractions. Immunofluorescence microscopy of non-purified peripheral blood mononuclear cell (PBMC) fractions treated with the NET stimuli, PMA (50 nM, 2.5h) or nigericin (15 µM, 2.5 h), and then stained with Hoechst, anti-neutrophil elastase (NE) and 3D9 or PL2.3. Images were taken on an upright fluorescent microscope at 20X magnification. The selected images are representative of 3 independent experiments. Scale bar – 50 µm.

### 3D9 distinguishes NETosis from other forms of cell death in neutrophils

Neutrophils can commit to other cell death pathways (recently reviewed by Dabrowska et al., 2019) besides NETosis. Naïve neutrophils undergo apoptosis after overnight incubation (Kobayashi et al., 2005) and necroptosis upon TNFα stimulation in the presence of a SMAC (second mitochondria-derived activator of caspase) mimetic and if caspases are inhibited (Galluzzi et al., 2012). Interestingly, the anti-chromatin antibody (Figure 8-figure supplement 1), but not 3D9, stained the condensed nuclei of cells undergoing apoptosis (Figure 8) and neither of the antibodies stained cells during necrosis induced by the staphylococcal toxin α-haemolysin nor after stimulation with necroptosis inducers.

**Figure 8.**
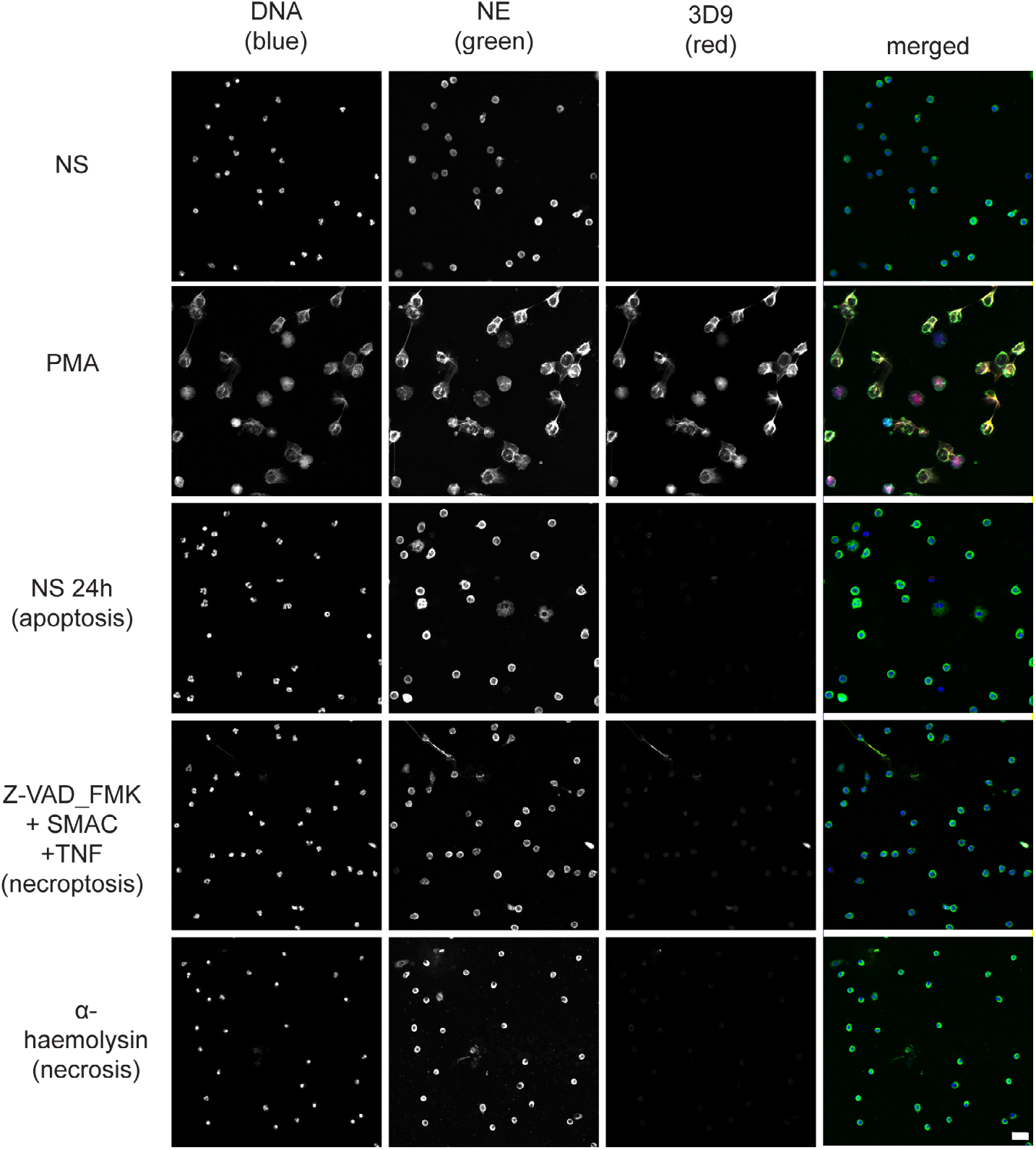
Comparison of 3D9 detection in response to apoptotic, necroptotic & necrotic cell death stimuli. Confocal immunofluorescent microscopy of neutrophils stimulated with different cell death stimuli and subsequently stained with Hoechst, anti-neutrophil elastase (NE) and 3D9. NETs were induced with PMA (100 nM, 3h). Apoptosis was induced in resting neutrophils by incubation for 24h without stimulation overnight. Neutrophils were stimulated with Z-VAD-FMK (50 µM) plus SMAC mimetic (100 nM) plus TNF (50 ng/ml) for 6h to induce necroptosis. Necrosis was induced with the pore forming toxin α-haemolysin (25 µg/ml). Images were taken at 20X and are representative of 3 experiments. Scalebar 20 µm. A comparison was made with parallel samples stained with the chromatin antibody PL2.3 and are presented in Figure 8-figure supplement 1.

### 3D9 labels NETs in human tissue sections

NETs are found in inflamed tissues based on the juxtaposition of chromatin and granular makers as well as the detection of citrullinated H3. 3D9 stains areas of decondensed DNA (Hoechst) that colocalise with NE in both inflamed human tonsil (Figure 9A) and human kidney (Figure 9B). This indicates that 3D9 labels NETs in histological samples - hematoxylin and eosin (HE) tissue overviews are provided in the supplemental figures (Figure 9-figure supplement 1; Figure 10-figure supplement 1; Figure 11-figure supplement 1). Indeed, in kidney (Figure 9C), inflamed appendix (Figure 10) and gallbladder (Figure 11), 3D9 labelled DNA in the same cluster as anti-H3cit or anti-H2B. Interestingly, 3D9 stained decondensed, more NET-like structures, while anti-H3cit or anti-H2B antibodies stained relatively compact chromatin. Furthermore, colocalization analysis of 3D9 with H2B or with H3cit revealed that 3D9 was more commonly colocalised with H2B as compared to H3cit; overlap coefficients 0.463 (3D9-H2B) v 0.125 (3D9-H3cit), and 0.533 (3D9-H2B) v 0.122 (3D9-H3cit) for Figure 10. figure supplement 1 and Figure 11-figure supplement 1 respectively.

**Figure 9.**
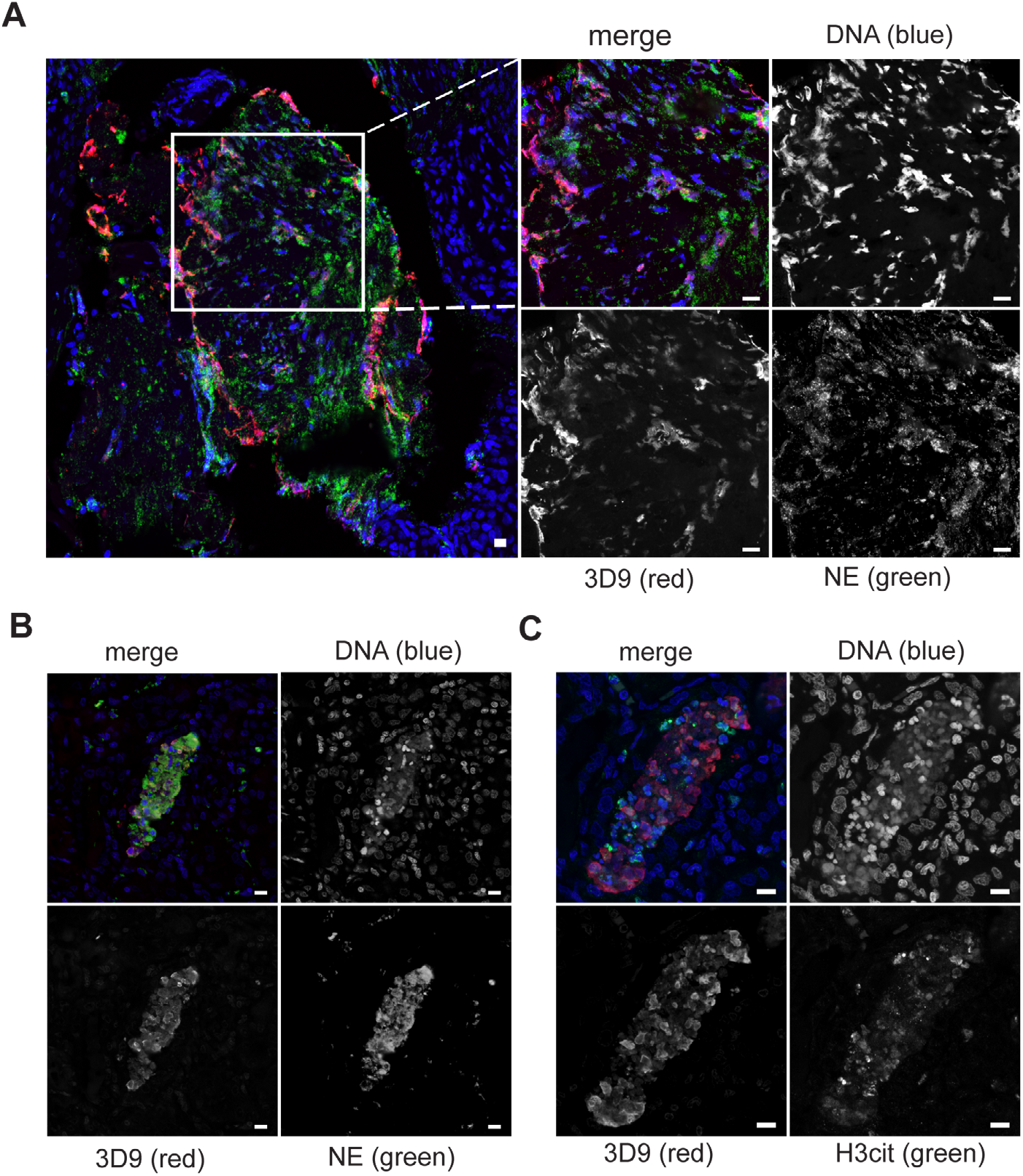
Detection of clipped histone 3 & NETs in human tissues. Paraffin embedded sections were stained with Hoechst, anti-NE and 3D9 or H3cit antibodies and examined by confocal microscopy. Scale bar - 10 µm **(A**) inflamed human tonsil (**B** & **C**) Inflamed human kidney.

**Figure 10.**
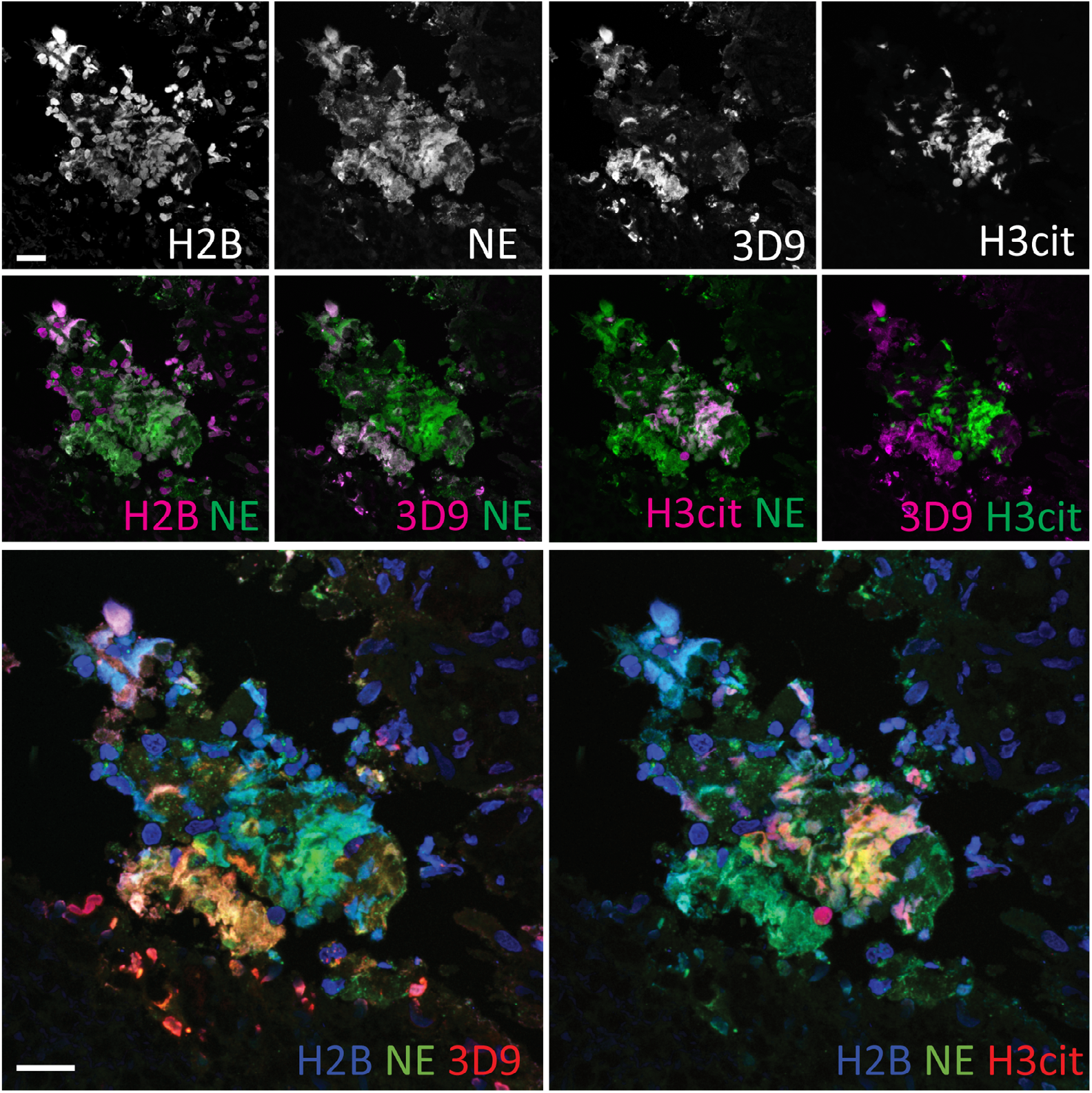
Comparison of Clipped H3, H3cit & H2B staining in the gallbladder. Paraffin embedded sections were stained with Hoechst, anti-NE and histone antibodies 3D9, H3cit and H2B and examined by confocal microscopy. Scale bar – 20 µm.

**Figure 11.**
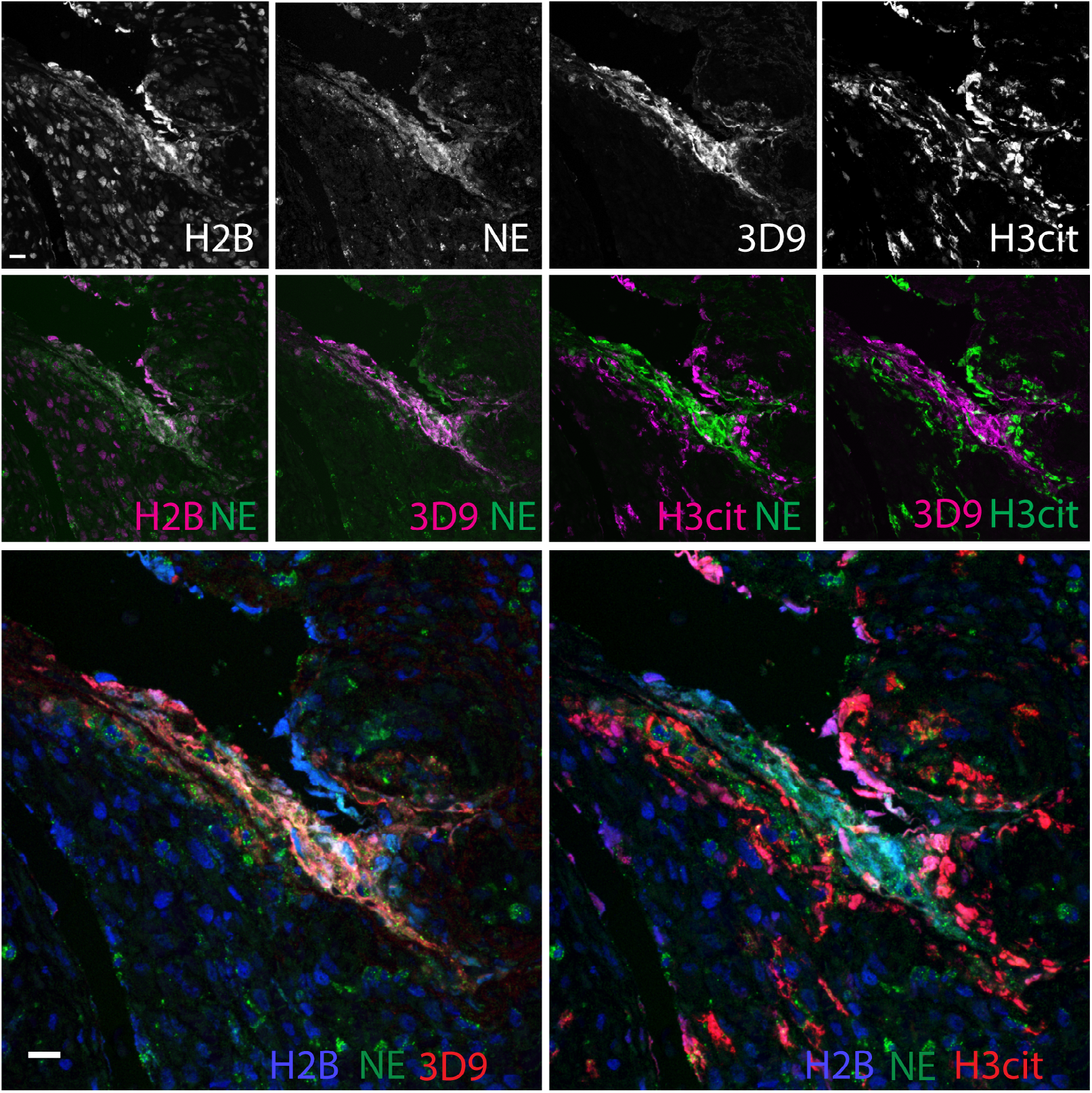
Comparison of Clipped H3, H3cit & H2B staining in the appendix. Paraffin embedded sections were stained with Hoechst, anti-NE and 3D9 or H3cit antibodies and examined by confocal microscopy. Scale bar - 20 µm.

## Discussion

Decondensed chromatin is a defining feature of NETs. It occurs through PTMs that partially neutralise the histone positive charge and thus the affinity of histones for negatively charged DNA (Papayannopoulos et al., 2010; Wang et al., 2009). One way to achieve this is through proteolytic removal of the lysine and arginine rich histone tails. Using a biochemical and proteomic approach, we determined that H3 is cleaved within its globular domain during NETosis. We exploited the specificity of this event to produce a mouse monoclonal antibody to the *de novo* histone H3 epitope, the new N-terminal beginning at R49. This antibody, 3D9, recognises human NETs induced by both microbial and host derived physiological stimuli and distinguishes netotic neutrophils from neutrophils that die via alternative pathways. It also discriminates between NETs and other cells in mixed blood cell fractions and, importantly, NETs in human histological samples.

Until now, histone citrullination is the only PTM that has been used for antibody-based detection of NETs. In this paper, we propose histone cleavage at H3R49 as a new histone PTM that can be used for broad detection of NETs from human samples. The use of H3cit for the detection of NETs is not without controversy. Not all NETs are citrullinated and NET formation can occur in the absence or inhibition of citrullinating enzymes (Kenny et al., 2017; Konig and Andrade, 2016). By identifying the precise histone cleavage site, we shed further light on this. The most commonly used H3cit antibody detects citrullination of R2, R8 and R17. However, histone cleavage at R49 would remove the H3cit epitopes, rendering these NET defining PTMs mutually exclusive on a single histone level. Indeed, co-staining by anti-H3cit and 3D9 in inflamed kidney, gallbladder and appendix paraffin sections revealed extensive mutual exclusion of the two marks and more abundant staining of decondensed chromatin by 3D9. Thus, we propose that 3D9 will allow broad detection of NETs but may display a preference for more mature or proteolytically processed NETs.

In contrast to the present study, histone cleavage was reported as discriminating between different pathways of NET formation (Pieterse et al., 2018). Using a sandwich ELISA approach, Pieterse *et al* concluded that, generally, the N-terminal histone tails are removed in NOX dependent but not NOX independent NET formation. They used a suite of N-terminal directed antibodies for H2B, H3 and H4, and all H3 epitopes were located N-terminal to the cleavage site H3R49. However, in the final biological sample testing the authors did not examine H3. Interestingly, the authors observed that, by immunofluorescent microscopy, all histone N-terminal antibodies failed to stain NETs at time points after cell lysis. Therefore, to us, this data suggests that at later stages in NETosis the histone N-terminal tails, at least for H3, are removed irrespective of the pathway of activation. This is in line with our observations that both NOX dependent (PMA, heme) and NOX independent (nigericin) stimuli result in NETs that are recognised by 3D9 and supports our finding that histone H3 cleavage at R49 is a general feature of human NET formation.

In this study we detect NETs in fixed or denatured human samples from *in vitro* experiments and histological samples. While an ELISA with 3D9 revealed a preference for NETs over isolated chromatin or recombinant H3, immunoprecipitation of naïve cell lysates by 3D9 also pulled down full length H3 as confirmed by immunoblot. It is not yet clear if this is due to co-immunoprecipitation due to the presence of low levels of clipped histone or if this represents true recognition of intact H3 by 3D9. Thus, care should be taken when detecting cleaved H3 or NETs under native and mild detergent conditions and all sample types e.g. serum samples, need careful validation for cross reactivity.

More broadly, and applying to the general principles of NET detection, it is not yet possible to prove conclusively that the detected decondensed chromatin originates from the same cell source as the neutrophil proteins which decorate it. For example, in an infected necrotic wound to which to high numbers of neutrophils are recruited. Here, activated neutrophils might release both proteases (Borregaard et al., 1993) and citrullinating enzymes (Spengler et al., 2015; Zhou et al., 2017) that bind to and modify extracellular chromatin generating a NET - according to the histological definition. This remains a conundrum that requires further exploration.

N-terminal histone cleavage at H3R49 is a novel and so far undescribed cleavage site in any eukaryotic organism. Unusually, it is located in the globular rather than the unstructured tail region of H3. H3R49 is one of 6 key residues important for the regulation of H3K36me^3^ and forms part of the structured nucleosome surface (Endo et al., 2012). Thus, we speculate that removal of the N-terminal tail, in its entirety, could lead to removal of higher order structure interactions and facilitate chromatin decondensation e.g. removal of H3K9me and its associated heterochromatin protein 1 interactions (Jacobs and Khorasanizadeh, 2002). To determine the contribution of histone cleavage at H3R49 to the process of chromatin decondensation, future work will focus on establishing the protease(s) responsible and the sequence of proteolytic events leading to this final truncation of H3 and NET formation. Given the specificity of this event and its restriction to NETotic forms of cell death, we propose that N-terminal cleavage at H3R49 is an example of histone ‘clipping’ in neutrophils – a term proposed by the histone/histone proteolysis field for specific histone cleavage sites for which a biological function has been demonstrated (Dhaenens et al., 2015).

In conclusion, this study represents the first identification of a distinctive and exclusive marker of NETs and describes the development and characterisation of a complementary antibody to facilitate easier detection of human NETs. Analogous to finding a smoking gun at a crime scene, the monoclonal antibody 3D9 detects evidence of the proteolytic events that occur in NETosis – the proteolytic signature, histone cleavage at H3R49. In doing so, 3D9 discriminates NETs from chromatin of other cells and chromatin of neutrophils that die via alternative mechanisms. This added layer of specificity will simplify the detection of NETs in tissue samples and facilitate comparison of quantitative studies between labs. This will be an important step in assessing the contribution of extracellular chromatin and NETs to disease pathology.

## Materials and methods

### Reagents

All reagents were purchased from common vendors of laboratory reagents e.g. Sigma Aldrich or VWR Deutschland unless otherwise stated.

### Blood collection and ethical approval

Venous blood was collected from healthy donors who had provided informed consent according to the Declaration of Helsinki. Ethical approval was provided by the ethics committee of Charité-Universitätsmedizin Berlin and blood was donated anonymously at Charité Hospital Berlin.

### Purification and culture of human peripheral blood neutrophils

Neutrophils were isolated as described by Amulic et al (2017). Briefly, venous whole blood was collected in EDTA and separated by layering over equal volume Histopaque 1119 and centrifugation at 800 *g* (20 min). The pinkish neutrophil rich fraction was collected and washed once by the addition of 3 volumes of wash buffer (PBS, without Mg^2+^ or Ca^2+^ [Gibco] supplemented with 0.5% [w/v] human serum albumin [HSA, Grifols]) and centrifugation at 300 *g* (10 min). The neutrophil fraction was further purified by density gradient centrifugation - Percoll (Pharmacia) gradient from 85%-65% (v/v). Purified cells were collected from the 80-70% fractions and washed once before being resuspended in wash buffer. Cells were counted using a CASY cell counter.

For all experiments, unless indicated, neutrophils were cultured RPMI (GIBCO 32404014) supplemented with 10 mM HEPES and 0.1% (w/v) HSA, which had been preequilibrated in CO_2_ conditions for 1 h. For some stimuli, the HSA content was reduced to 0.05% or 0% HSA as indicated in the figure legends. Cells were routinely cultured at 37°C, 5% CO_2_ unless indicated. For all experiments, stimuli were added to cell reactions as 10X working stock solutions freshly diluted in RPMI. For inhibition experiments, a 10X inhibitor stock and appropriate vehicle controls, were added to the cells and preincubated for the times stated in the figure legends.

### Neutrophil and NET lysate preparation

To analyse proteins, lysates were prepared from stimulated or resting neutrophils. Cells were seeded in culture medium in 1.5 ml microcentrifuge tubes at 1×10^7^ cells/ml with 5×10^6^ cells per time point. After addition of the inhibitor or agonist, cells were gently mixed and incubated at 37 °C with gentle rotation. At the specified time points, protease inhibitors - 1 mM AEBSF, 20 µM Cathepsin G inhibitor I (Calbiochem), 20 µM neutrophil elastase inhibitor GW311616A (Biomol), 2X Halt protease inhibitor cocktail (PIC, Thermofisher Scientific), 10 mM EDTA, 2 mM EGTA - were added directly to the cell suspension. Cells were gently mixed and centrifuged at 1000 *g* (30 s) to collect all residual liquid. Freshly boiled 5X sample loading buffer (50 mM Tris-HCl pH 6.8, 2% [w/v] SDS, 10% glycerol, 0.1% [w/v] bromophenol blue, 100 mM DTT) was added to samples which were then briefly vortexed and boiled (98 °C) for 10 min with agitation and flash frozen in liquid nitrogen for storage at −80 °C.

### 1D SDS-PAGE and immunoblot blot

For routine protein analysis, samples were analysed by 1D SDS-PAGE and immunoblotted. Samples were thawed on ice, boiled at 98 °C (10 min) and sonicated to reduce viscosity (Braun sonicator, 10 s, cycle 7, power 70%). Proteins were applied to NuPAGE 12% gels (Invitrogen, Thermofisher) and run at 150 V in MES buffer (Thermofisher Scientific). Proteins were transferred by western blot onto PVDF (0.2 µm pore size, Amersham GE Healthcare) using the BioRad wet transfer system (buffer: 25 mM Tris, 192 mM glycine, 20% methanol, protocol: 30 min at 100 mA, 120 min at 400 mA). Blotting efficacy was assessed by Ponceau S staining. Blots were blocked with TBST (TBS pH 7.5, 0.1% [v/v] Tween-20) with 5% [w/v] skimmed milk, for 1 h at RT. Blots were then incubated with the following primary antibodies overnight at 4 °C or for 2 h at RT: rabbit anti-histone H3 C-terminal pAb, 1:15000 (Active motif #61277); rat anti-histone H3 N-terminal mAb, 1:1000 (Active Motif #61647, aa 1-19); Histone H4 C-terminal, 1:5000 (Abcam 10158 – aa 50 to C terminal); rabbit anti-histone H4 N-terminal mAb, 1:30,000 (Upstate, Millipore #05-858, aa17-28); rabbit GAPDH mAb, 1:5000 (Cell Signalling Technology, #2118); mouse 3D9 1ug/ml (produced in this study) – all diluted in TBST with 3% (w/v) skimmed milk. After washing with TBST (3 x 5 min), blots were blocked for 15 min as before and then probed with secondary HRP conjugated antibodies (Jackson ImmunoResearch -diluted 1:20000 in 5% skimmed milk TBST) for 1 h at RT. Blots were washed in TBST (3 x 5 min) and developed using SuperSignal™ West Dura Extended Duration Substrate (ThermoFisher Scientific) and an ImageQuant Gel imager (GE Healthcare).

### Immunofluorescent staining of in vitro samples

For immunofluorescent imaging of purified cells, neutrophils/PBMCs were seeded in 24 well dishes containing glass coverslips, with 1×10^5^ cells per well and incubated at 37 °C for 1 h to allow to adhere to the coverslip. At this stage, inhibitors and priming factors were included as indicated. Reactions were stopped by the addition of paraformaldehyde (2% [w/v]) for 20 min at RT or 4°C overnight. After fixation, cells were washed and stained as previously described (Brinkmann et al 2012). Briefly, all steps were performed by floating inverted coverslips on drops of buffer on laboratory parafilm. Cells were permeabilised with PBS, pH 7.5, 0.5% (v/v) Triton X-100 for 3 min. For screening and quantification experiments, this permeabilization step was extended to 10 min. Samples were washed (3 x 5 min) with PBS, 0.05% (v/v) Tween 20 and incubated with blocking buffer - PBS pH 7.5, 0.05% (v/v) Tween 20, 3% (v/v) normal goat serum, 3% (w/v) freshwater fish gelatin, 1% (w/v) BSA - for 20 min at RT and then probed with primary antibodies diluted in blocking buffer and incubated overnight at 4 °C. Primary antibodies: anti-chromatin (H2A-H2B-DNA complex) mouse mAb, 1 µg/ml (Losman 1992); neutrophil elastase, rabbit pAb 1:500 (Calbiochem); mouse serum for screening, 1:100; hybridoma supernatants, neat; 3D9 mouse mAb, 1 µg/ml. Samples were then washed as before. Alexa labelled secondary antibodies (Invitrogen) were diluted 1/500 in blocking buffer and incubated for 2 h at RT. DNA was stained with Hoescht 33342 (Invitrogen, Molecular Probes) 1 ug/ml, incubated with the secondary antibody step. Samples were washed in PBS followed by water and mounted in Mowiol mounting medium.

### Histone extraction from neutrophils

Histone enriched fractions were prepared from resting neutrophils and NETs according to a method modified from Shechter et al (2007). Neutrophils (4-8 ×10^7^) were resuspended in 13 ml of RPMI (without HSA) in a 15 ml polypropylene tube and incubated on a roller at 37 °C with PMA 50 nM for 90 min. After stimulation, 1 mM AEBSF was added to inhibit further degradation by NSPs and cells were cooled on ice for 10 min. All subsequent steps were performed on ice or 4 °C where possible. Cells and NETs were pelleted by centrifugation at 1000 *g*, 10 min. Samples were resuspended in ice-cold hypotonic lysis buffer (10 mM Tris-HCl pH 8.0, 1 mM KCL, 1.5 mM MgCl_2_, 1 mM DTT supplemented with protease inhibitors just before use – 1 mM AEBSF, 20 µM NEi, 20 µM CGi, 2X PIC, 10 mM EDTA) using 1 ml of buffer/5×10^6^ cells. Cells were then incubated at 4 °C on a rotator for 30 min before being passed through a syringe to aid lysis and shearing of intact cells. Nuclei and NETs were collected by centrifugation at 10,000 *g*, 10 min, discarding the supernatant. To disrupt nuclei, samples were resuspended in dH_2_O (1 ml/1×10^7^ cells) supplemented with protease inhibitors, as before, and incubated on ice for 5 min with intermittent vortexing. NP40 (0.2% [v/v]) was added to help lysis and disruption of NETs and samples were sonicated briefly (10 s, mode 7, power 70%). To extract histones, H_2_SO_4_ (0.4 N) was added to samples, and vortexed briefly. Samples were then incubated, rotating, for 2-3h. Histone enriched fractions were collected by aliquoting samples into multiple 1.5 ml microcentrifuge tubes and centrifuging at 16,000 *g* for 10 min, followed by collection of the supernatants. To minimise further processing of histones, proteins were immediately precipitated overnight by dropwise addition of trichloroacetic acid to a final concentration of 33% followed by mixing. The next day precipitated proteins were pelleted by centrifugation at 16,000 *g*, 10 min. The supernatants were discarded and waxy pellets were washed once with equal volume ice-cold acetone with 0.2% (v/v) HCl and 5 times with ice-cold acetone alone. Pellets were allowed to air dry for 5 min before being resuspended with 1 ml (per 5×10^6^ cells) of dH_2_O plus 1 mM AEBSF. For difficult to resuspend pellets the mixture was vigorously shaken at 4°C overnight before samples were centrifuged, as before, to remove undissolved protein. Pooled supernatants for each sample were lyophilised and stored at −80°C until histone fractionation.

### Purification of histone H3

Histones were fractionated by RP-HPLC according to the method described by Shechter et al (2007). After lyophilisation samples were resuspended in 300 µl Buffer A (5% acetonitrile, 0.1% trifluoroacetic acid) and centrifuged at 14,000 *g* to remove particulate matter. 150 µl of clarified sample was mixed with 40 µl Buffer A before being applied to a C18 column (#218TP53, Grave Vydac) and subjected to RP-HPLC (Waters 626 LC System, MA, US) as described by Schecter et al (2007). The flow rate was set to 1 ml min^-1^ and fractions were collected at 30 s intervals from minute 30 to 55. All fractions were lyophilised and stored at −80 °C until analysis. To determine which fractions contained H3 and cleaved species, each fraction was dissolved in 50 µl dH_2_O and 5 µl was subjected to SDS-PAGE and either stained with Coomassie blue stain or transferred to PVDF and immunoblotted for H3 and H4 as described already.

### Two dimensional electrophoresis (2-DE) of purified histones

To determine the cleavage sites, H3 containing fractions were pooled and subjected to a small gel 2-DE procedure (Jungblut and Seifert, 1990). Briefly, pooled fractions were denatured in 9 M urea, 70 mM DTT, 2% Servalyte 2-4 and 2 % Chaps. Samples (30 µl) were applied to 1.5 mm thick isoelectric focusing (IEF) gels using ampholytes 7-9 and a shortened IEF protocol was used: 20 min 100 V, 20 min 200 V, 20 min 400 V, 15 min 600 V, 5 min 800 V, and 3 min 1000 V, (a total of 83 min and 500 Vh) in 8 cm long IEF tube gels. Separation in the second dimension was performed in 6.5 cm x 8.5 cm x 1.5 mm SDS-PAGE gels. Duplicate gels were prepared; one stained with Coomassie Brilliant Blue R250 for excision of spots for mass spectrometry identification; and the second transferred to PVDF as follows. Proteins were blotted onto PVDF blotting membranes (0.2 µm) with a semidry blotting procedure (Jungblut et al., 1990) in a blotting buffer of 100 mM borate. Spots were stained by Coomassie Brilliant Blue R250 and analysed by N-terminal Edman degradation sequencing (Proteome Factory, Berlin, Germany).

### Antibody generation

Immunising and screening peptides are outlined in S.Table 2 (Figure 3-figure supplement 2) and were synthesized by Eurogentec (Belgium). A portion was further conjugated to Key Lymphocyte Haemoglutinin (KLH) for immunization. Immunisation of mice, preliminary ELISA screening and production of hybridomas were performed by Genscript as follows. Six mice (3x Balb/c and 3x C57/BL6) were immunized with branched peptides. Mice were bled and effective immunization was assessed using a direct ELISA. The ELISA and subsequent inhouse immunoblot and immunofluorescent microscopy screening strategy are outline in Figure 3-figure supplement 1. Following selection of effectively immunized animals, a further boost injection of the immunogen was given before isolation of spleen cells for hybridoma production. The resulting hybridoma supernatants were screened similarly. Large-scale culture of supernatants and purification of antibodies was performed by Genscript.

### ELISA

To assess 3D9 specificity for NETs, 3D9 was used in an indirect ELISA to detect cleaved H3 in purified NETs, chromatin, recombinant H3 and DNA. NETs were prepared by seeding 3×10^6^ neutrophils in a 6 well dish and incubating for 3-6 h with 100 nM PMA. NETs were gently washed 3 times with equal volume PBS, before being collected in 300 µl PBS. Clumped NETs were disrupted by sonicating briefly (3 s, mode 7, power 70% - Braun Sonicator) and then snap frozen and stored at −80°C. Chromatin was prepared from lung epithelial cells (A549) as previously described by Shechter *et al* (2007). The final nuclear pellet was resuspended in dH_2_O and sonicated as before and stored at −80 °C. The DNA content of NETs and chromatin was assessed by PicoGreen assay according to the manufacturer’s instructions (Thermofisher Scientific). Beginning at 1 µg/ml (of DNA content), serial dilutions of NETs, chromatin and calf thymus DNA (Invitrogen) were prepared in lo-DNA bind Eppendorf microcentrifuge tubes. A similar dilution series of recombinant histone H3 (New England Biolabs) was prepared starting at 1 µg/ml protein. All dilutions were performed in PBS. One hundred microliters of each sample, in duplicate, at dilutions 1 ug/ml to 1 ng/ml, was aliquoted in a Nunc Maxisorb 96 well dish and immobilised overnight at 4°C, 250 rpm. The immunising peptide, REIRR (10ng/ml) was used as a positive control. The following day all wells were washed 6 times with wash buffer (PBS, 0.05% Tween 20) and then blocked with 200 µl of blocking solution (1% BSA in wash buffer) for 2 h (RT). Wells were washed once with wash buffer and incubated with 100 µl of 3D9 (2 µg/ml, in blocking solution) at RT (2 h) with gentle shaking (250 rpm). Wells were washed 6 times as before and then incubated with 100 µl of secondary HRP conjugated anti-mouse (Jackson laboratories) at 1:10,0000 dilution in blocking solution and incubated as before. Finally, wells were washed 6 times as before and HRP activity was detected using TMB (3,3’, 5,5’ tetramethylbenzidine) reagent (BD OptEIA™) according to the manufacturer’s instructions (incubating for 15 −30 min). The assay was stopped by the addition of 100 µl H_2_SO_4_ (0.16 M) and absorbance (450 nm) was measured on a 96 well plate reader (VERSAmax, Molecular Devices, CA, US).

### Quantification of staining characteristics by Image J and R

In order to assess staining characteristics of antibodies during NET formation we developed a bundle of Image J and R scripts. These scripts allow for an automated workflow starting from 2-channel microscopic images of an experimental series (DNA stain, antibody stain), to a graphical representation and classification of individual cells and eventually to mapping these classifications back to the original images as a graphical overlay. In the first step nuclei are segmented based on intensity thresholding (either programmatic or manual), including options for lower and upper size selection limits. The same threshold is applied to the entire experimental series and the upper size limit is used to exclude fused structures that cannot be assigned individual cells. A quality score is assigned to every image based on the fraction of the total DNA stained area that can be assigned to individual cells (or NETs). This score along with all other parameters of the analysis is exported as report file and can be used to automatically exclude images from the analysis. In addition, this part of the script generates a result file that includes the area, circumference (as x,y coordinates) and cumulative intensities for each channel for every detected nucleus along with information such as time point or stimulus that can be assigned programmatically. In order to analyse these data sets we implemented a series of functions in R. These functions include import of Image J result files, classification of cells based on nuclear area and staining intensity, various plot and data export functions, as well mapping functions that allow to display the classification of nuclei as color coded circumferences overlaid on the original images. The scripts are available for download at https://github.com/tulduro/NETalyser

### Quantification of NETs by Image J

NETs were quantified by the semi-automatic method described by Brinkmann et al (2012) and via a second modified method that allowed automatic quantification. All microscopy image datasets were processed by both methods to allow comparison. Hoechst was used to stain total DNA and NETs were additionally stained with anti-chromatin (PL2.3) or 3D9 antibodies (1 µg/ml) and Alexa-568 coupled secondary antibody according to the previous section. Images were acquired with a Leica DMR upright fluorescence microscope equipped with a Jenoptic B/W digital microscope camera and 10x or 20x objective lens. For each experiment, the same exposure settings were used for all samples and a minimum of 3 random fields of view (FOV) were collected. Images were analysed using ImageJ/FIJI software. As described by Brinkmann et al (2012), each channel was imported as an image sequence and converted into a stack. To count total cells/NETs per FOV, the Hoechst channel stack was imported and segmented using the automatic thresholding function (Bernsen method) with radius 15 and parameter 1 set to 35 to produce a black and white thresholded image. Particle analysis was then performed to count all objects the size of a cell nucleus or bigger (10x objective: particle size 25-infinity; 20x objective: particle size 100-infinity) and to exclude background staining artefacts. Total NETs were then counted using the anti-chromatin (PL2.3) or 3D9 channels accordingly. For the method published in 2012, anti-chromatin stains were segmented using manually adjusted thresholding so that the less intense staining of resting cell nuclei was excluded. Particle analysis was then performed to count all objects larger than a resting cell nucleus (10x objective: particle size 75-infinity; 20x objective: particle size 250-infinity). This was also performed for 3D9. In contrast, in the second analysis, the workflow was modified so that the automatic Bernsen thresholding and segmentation were used for both Hoechst and NET channels, total cells and NETs respectively. For each method percentage NETs were calculated as (NETs/Total cells) x100. Results per FOV were then averaged according to sample. A schematic of the different workflows is presented in.Figure 5-supplement 1.

### Immunofluorescent staining of histological samples from tissue sections

Paraffin sections (2 µm thick) were deparaffinized in two changes of 100% xylene for 5 min each and then rehydrated in two changes of 100% ethanol for 5 min each and followed by 90% and 70% ethanol for 5 min each. Sections were washed with 3 changes of deionized water and incubation in TBS (Tris buffered saline). For antigen retrieval, Target Retrieval Solution (TRS pH9) (Dako S2367) was used to incubate the slides in a steam cooker (Braun) for 20 min. After cooling down to room temperature in antigen retrieval buffer, slides were rinsed 3x in deionized water and incubated in TBS until further processing. Slides were blocked with blocking buffer (1% BSA, 5% normal donkey serum, 5% cold water fish gelatin and 0.05% Tween20 in TBS, pH7.4) for 30 min. Blocking buffer was removed, and sections were incubated with primary antibodies at appropriate dilution in blocking buffer (containing 0.05% Triton-X100) overnight at RT. Sections were rinsed in TBS and then incubated with secondary antibody at an appropriate concentration (1:100) for 45 min in the dark at RT and then rinsed three times in TBS for 5 min followed by rinsing with deionized water. Slides were incubated with DNA stain Hoechst 33342 (1:5000) for 5 min, rinsed with water before mounting with Mowiol. Primary antibodies for detection of NETs were as follows: mouse anti-cleaved histone 3 clone 3D9 (2 µg/ml); rabbit anti-histone H3 antibody (citrulline R2 + R8 + R17; ab5103, Abcam); chicken anti-Histone H2B ab134211 Abcam (1:400); sheep anti-ELANE (NE) LS-Bio LSB 4244 Lot 75251. Secondary antibodies were the following: donkey anti-rabbit immunoglobulin G (IgG) heavy and light chain (H&L) Alexa Fluor 488 (Jackson 711-225-152); and donkey anti-mouse IgG H&L Cy3 (Jackson 715-165-151); donkey anti-sheep (IgG) H&L Alexa Fluor 647 (Jackson 713-605-147). An upright widefield microscope (Leica DMR) equipped with a JENOPTIK B/W digital microscope camera or a Leica confocal microscope SP8 were used for fluorescent imaging. Z-stack images were collected at 63× magnification. Where stated colocalisation analysis was performed on confocal images using Volocity 6.5.1 software. Human tonsil and kidney paraffin tissue blocks were purchased from AMSbio. Inflamed tissue from a gallbladder and an appendix was obtained from archived leftover paraffin embedded diagnostic samples and used in an anonymised way after approval through the Charité Ethics Committee (Project EA4/124/19, July 24, 2019). Informed consent from patients for use of biomaterials for research was obtained as part of the institutional treatment contract at Charité.

### Statistics

All experiments were repeated 3 times unless stated differently in the figure legend. Experimental repeats are biological replicates, where each replicate represents cells isolated from a different donor. All graphs were prepared in GraphPad Prism and are either representative traces or mean ± standard deviation as stated in the figure legend. Graphs for epitope mapping were produced by Pepscan using proprietary software.

## Supporting information

Supplemental File 1_epitope mapping peptides

## Contributions

DOT-Conceptualization, methodology, investigation, data curation, figure preparation, writing original draft, review and editing; UZA - 2-DE, blotting and sample preparation for protein sequencing; UA -preparation of human pathology tissue samples and microscopy of tissue samples, image curation; MS - MALDI mass spectrometry; SF - provision and description of human pathology tissues and advice on histology figure presentation; PRJ - proteomic and 2-DE analysis; VB - colocalisation analysis, image curation, reviewing and critical feedback on manuscript; AH - conceptualization, data curation for R analysis, R scripting; AZ-conceptualization, reviewing and editing of manuscript; AZ and AH provided critical feedback to DOT and helped shape the research, analysis and manuscript.

## Acknowledgements

The authors would like to give special thanks to Borko Amulic and Gerben Marsman for their constructive feedback at the early stages of manuscript preparation. This work was supported by the Max Planck Society.

## List of supplemental figures (located after references)

**Figure 1-figure supplement 1**. Cysteine protease inhibition by E64 does not inhibit histone H3 cleavage.

**Figure 1-figure supplement 2**. AEBSF does not inhibit ROS production

**Figure 1-figure supplement 3**. AEBSF is not cytotoxic.

**Figure 2-figure supplement 1**. Schematic summary of extraction & identification of histone H3 cleavage sites in NETs.

**Figure 2-figure supplement 2**. Separation of histone H3 by two dimensional electrophoresis

**Figure 2-figure supplement 3**. S.Table 1 : Mass spectrometry identification of proteins co-separating with histone H3 following 2-DE.

**Figure 3-figure supplement 1**. Outline of immunisation & screening strategy for antibody production.

**Figure 3-figure supplement 2**. S.Table 2: List of immunisation, screening and competition peptides.

**Figure 3-figure supplement 3**. Peptide inhibition of 3D9 binding to NETs

**Figure 3-figure supplement 4**. 3D9 immunoprecipitation

**Figure 4-figure supplement 1**. Binding profiles recorded for 3D9 on the linear peptide array.

**Figure 4-figure supplement 2**. Binding profiles recorded for 3D9 on helical peptide mimics arrays.

**Figure 4-figure supplement 3**. S.Table 3. Summary of identified 3D9 binding regions in the peptide array.

**Figure 4-figure supplement 4**. Fine epitope mapping by replacement analysis.

**Figure 5-figure supplement 1**. Workflow of NET analysis methods

**Figure 8-figure supplement 1**. Comparison of PL2.3 detection in response to apoptotic, necroptotic & necrotic cell death stimuli.

**Figure 9-figure supplement 1**. Hematoxylin & eosin (HE) stain of kidney section

**Figure 10-figure supplement 1**. Hematoxylin & eosin (HE) stain of gallbladder & colocalization analysis

**Figure 11-figure supplement 1**. Hematoxylin & eosin stain of inflamed appendix & colocalization analysis.

## List of supplemental files

**Supplementary file 1**. Peptides for epitope mapping

**Supplementary file 2**. Supplemental Methods

**Figure 1-Figure supplement 2-Source Data 1**. AEBSF does not inhibit the ROS burst

**Figure 1-Figure Supplement 3_Source Data 1**. AEBSF is not cytotoxic

**Figure 3-Source Data 1**. Direct ELISA for cleaved H3

**Figure 4-Figure supplement 1-Source Data** 1. Linear and helical peptide epitope mapping

**Figure 4-Figure supplement 2-Source Data 1**. Linear and helical peptide epitope mapping

**Figure 4-Figure supplement 4-Source Data 1**-Fine epitope mapping by replacement analysis

**Figure 5-Source Data 1**. Comparison of NET quantification using manual or automatic thresholding and segmentation procedures for chromatin antibody and 3D9

