## Supplementary file 3-Figure supplements for "Histone H3 clipping is a novel signature of human neutrophil extracellular traps"

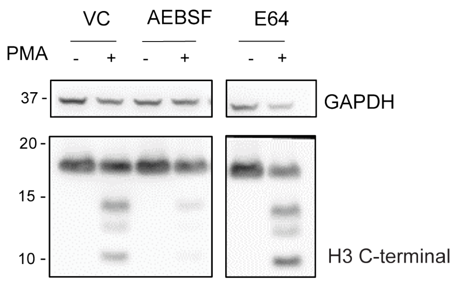


**Figure 1–figure supplement 1. Cysteine protease inhibition by E64 does not inhibit histone H3 cleavage.**
Neutrophils were preincubated with the cysteine protease inhibitor E64 (10 µM) or serine protease inhibitor AEBSF (150 µm) for 30 min and then stimulated with or without PMA 50 nM for 120 min. Lysates were resolved by SDS-PAGE and immunoblotted with anti-H3 C-terminal. GAPDH was used as a loading control. VC - vehicle control. Blot is representative of 3 independent experiments.

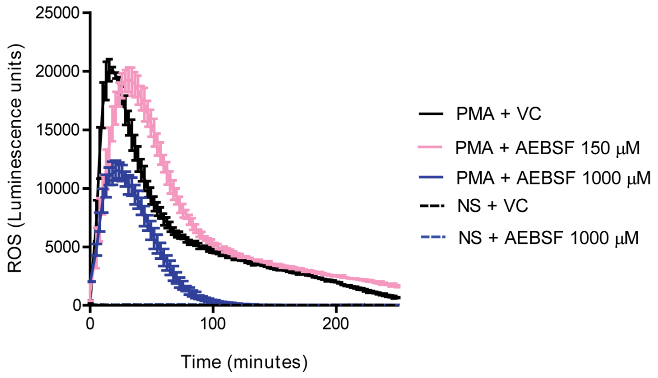


**Figure 1–figure supplement 2. AEBSF does not inhibit ROS production**
Neutrophils were preincubated with AEBSF at the specified concentrations for 30 min before stimulation as indicated. ROS production was assessed using a luminol-HRP assay. Representative trace of 3 independent experiments. VC – vehicle control; NS – non stimulated (PBS only). Source data can be found in Figure 1-figure supplement 2-Source Data 1.

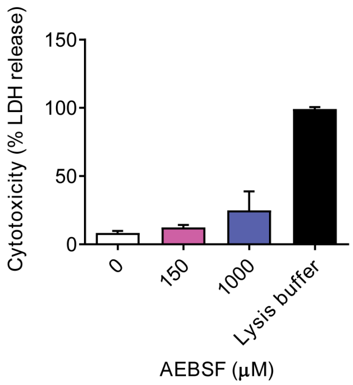


**Figure 1–figure supplement 3. AEBSF is not cytotoxic.**
Neutrophils were incubated with AEBSF at the indicated concentration and cytotoxicity was measured by LDH (lactate dehydrogenase) release where 100% lysis was achieved by adding 0.1 % (w/v) Triton X-100. Samples were measured at 490 nm and normalised to 100% lysis. Graph represents the mean ± standard deviation of 3 independent experiments. Source data can be found in Figure 1-figure supplement 3-Source Data 1.

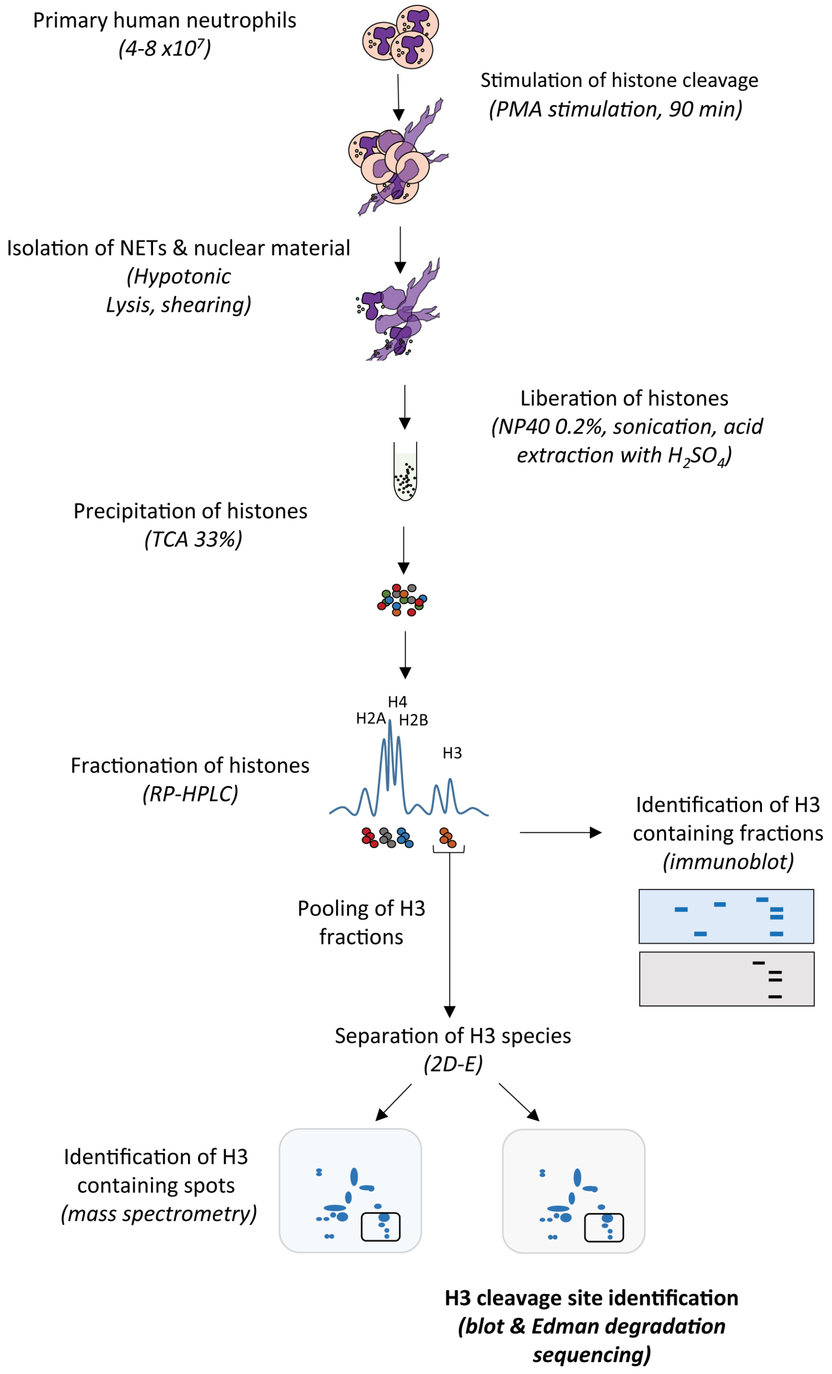


**Figure 2-figure supplement 1. Schematic summary of extraction & identification of histone H3 cleavage sites in NETs.**
Neutrophils were stimulated with PMA (50 nM) to induce histone cleavage and the samples were collected in the presence of proteases inhibitors. The nuclear material was isolated by hypotonic lysis and mild shearing. Nuclear material, including intact nuclei, was resuspended in $dH_2O$ plus protease inhibitors and NP40, and followed by gentle sonication to disrupt NETs. Histones were liberated from DNA by the addition of $H_2SO_4$ (0.4 M) and then precipitated with trichloroacetic acid (TCA) and solubilised in $dH_2O$ before separation by RP-HPLC on an acetonitrile gradient. Histone H3 containing fractions were identified by 1D-SDS-PAGE and immunoblotting for H3. H3 containing fractions were pooled, then divided in two, and both separated by 2D-electrophoresis (2D-E). Subsequent gels were either stained with Coomassie Blue and the

spots prepared for identification by mass spectrometry, or transferred to PVDF membranes, stained with Coomassie Blue and subjected to N-terminal Edman degradation sequencing.

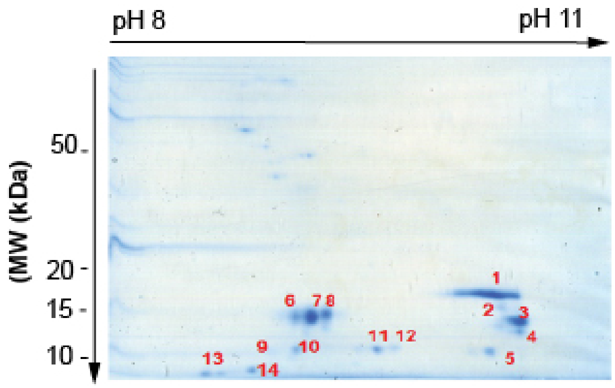


**Figure 2-figure supplement 2. Separation of histone H3 by two dimensional electrophoresis**
Coomassie stained gel of pooled histone H3 containing fractions resolved by two dimensional electrophoresis (2-DE). All labelled spots were subsequently analysed by mass spectrometry to identify the proteins in Figure 2-figure supplement 3.

***S.Table 1.*** *Mass spectrometry identification of proteins co-separating with histone H3 following RP-HPLC and 2-DE*

| spot | score | accession no. | protein name | MW | pI | sequence coverage | No. of peptides |
|---|---|---|---|---|---|---|---|
| **1** | 99 | P84243 | **Histone H3.3 (*H3*)** | 15318 | 11,27 | 25,7 | 8 |
| **2** | 133 | P84243 | **Histone H3.3 (*H3*)** | 15318 | 11,27 | 25 | 7 |
| **2** | 78 | P0C0S5* | **Histone H2A.Z** | 13545 | 10,58 | 14,1 | 4 |
| **3** | 140 | P84243 | **Histone H3.3 (*H3*)** | 15318 | 11,27 | 22,1 | 7 |
| **4** | 193 | P84243 | **Histone H3.3 (*H3*)** | 15318 | 11,27 | 27,2 | 8 |
| **5** | 198 | P84243 | **Histone H3.3 (*H3*)** | 15318 | 11,27 | 27,2 | 9 |
| **6** | 317 | P06702 | **Protein S100-A9** | 13234 | 5,71 | 70,2 | 10 |
| **7** | 486 | P06702 | **Protein S100-A9** | 13234 | 5,71 | 81,6 | 15 |
| **8** | 483 | P06702 | **Protein S100-A9** | 13234 | 5,71 | 82,5 | 14 |
| **9** | 57 | P31949* | **Protein S100-A11** | 11733 | 6,56 | 9,5 | 2 |
| **10** | 344 | P31949 | **Protein S100-A11** | 11733 | 6,56 | 45,7 | 8 |
| **11** | 343 | P05109 | **Protein S100-A8** | 10828 | 6,51 | 37,6 | 9 |
| **12** | 140 | P05109 | **Protein S100-A8** | 10828 | 6,51 | 25,8 | 5 |
| **13** | 136 | P25815 | **Protein S100-P** | 10393 | 4,75 | 35,8 | 6 |
| **14** | 48 | P06703* | **Protein S100-A6** | 10173 | 5,33 | 16,7 | 3 |

*candidate only

**Figure 2-figure supplement 3. S.Table 1**: Mass spectrometry identification of proteins co-separating with histone H3 following 2-DE. The top identifications are listed.. Due to the organization of the database, H3.3 is the top hit although the sequence coverage of the identified peptides does not allow distinction of histone H3 variants.

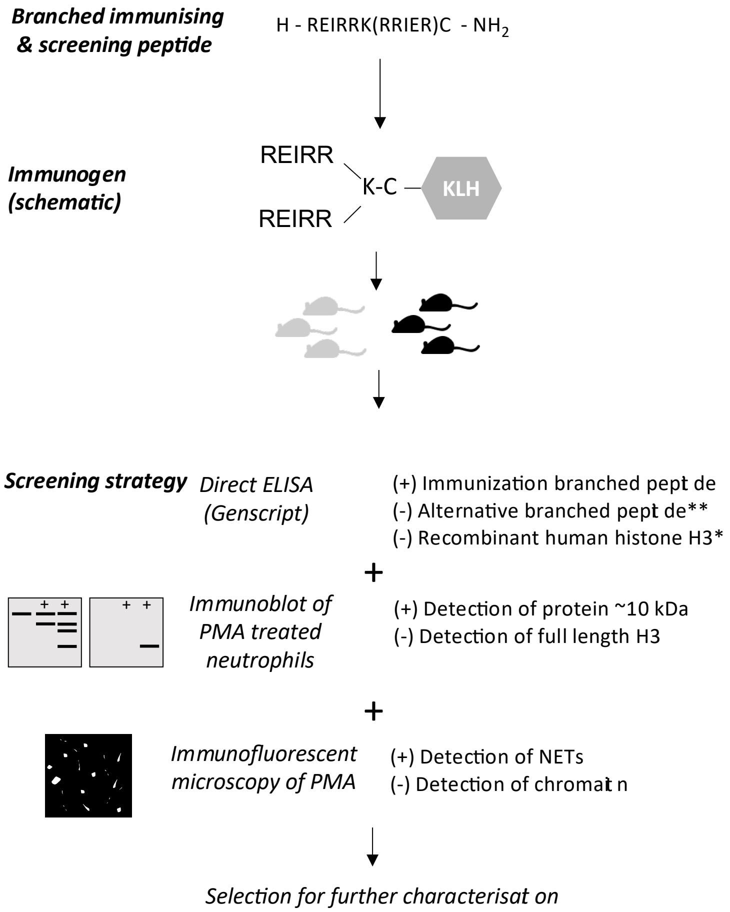


**Figure 3-figure supplement 1. Outline of immunisation & screening strategy for antibody production.** After selection of animals for hybridoma production, clone supernatants were screened similarly. Positive selection (+), negative selection (-). *Performed in the hybridoma screen only. ** Alternative branched peptide used for screening.

**S.Table 2. List of immunisation, screening and competition peptides**

| | |
|---|---|
| ***Immunisation*** | H - REIRRK(RRIER)C - $NH_2$ (KLH conjugated) |
| ***Screening*** | (+) H - REIRRK(RRIER)C -$NH_2$<br>(-) H - AARKSK(SKRAA)C-$NH_2$** |
| ***Validation*** | H -REIRRK(RRIER) - $NH_2$(competing peptide)<br>H -TGGVKK(KVGGT) - $NH_2$ (negative control) |

**Figure 3-figure supplement 2. S.Table 2: List of immunisation, screening and competition peptides.** Outline of the immunising peptides, the positive and negative screening peptides, and positive and negative control peptides for validation of the target by competition assay. The N-terminal is represented by H ($H_2N$) and the C-terminal is represented by $NH_2$ ($CONH_2$). ** Alternative branched peptide used for screening.

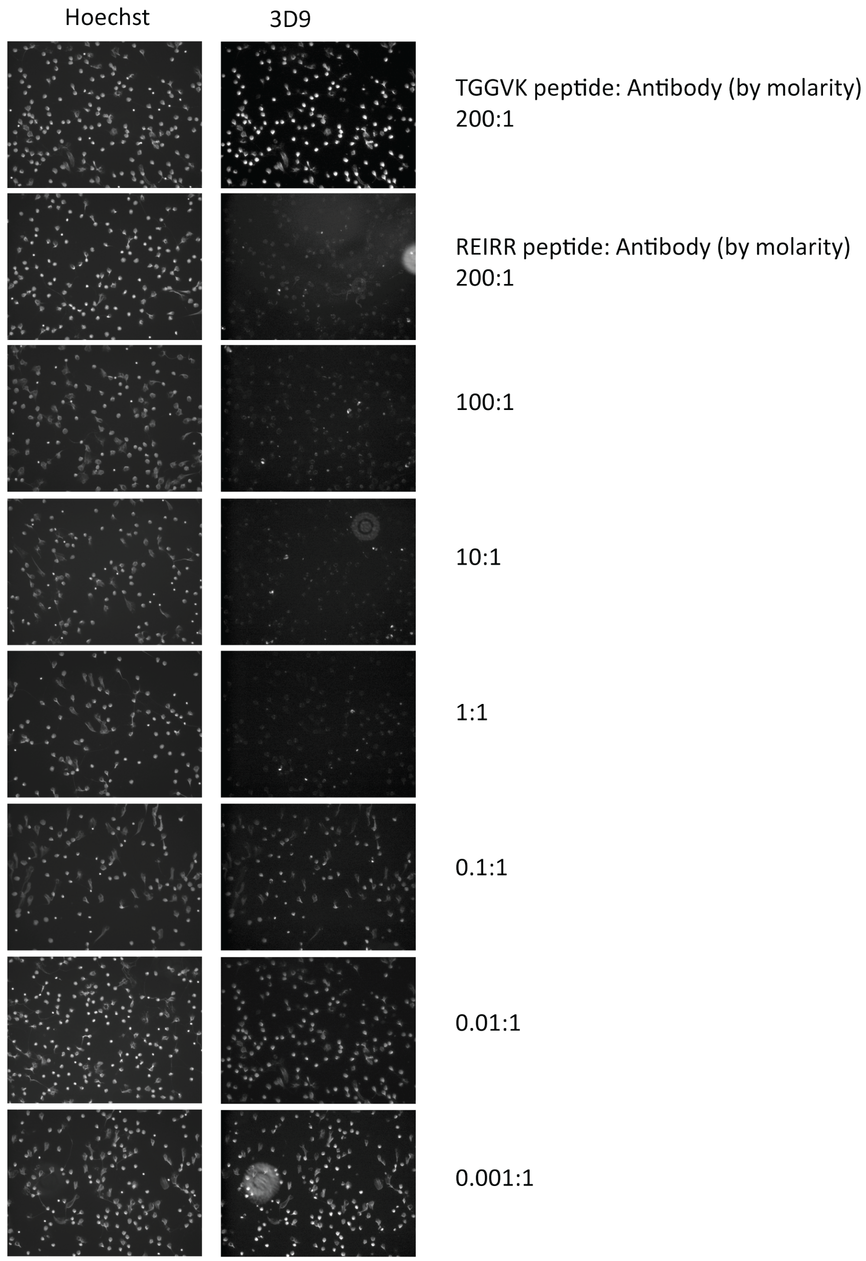


**Figure 3-figure supplement 3. Peptide inhibition of 3D9 binding to NETs**
Immunofluorescent microscopy of neutrophils stimulated with PMA for 3h and stained with 3D9 (1 ug/ml) in the presence of competition and control peptides. Prior to staining, 3D9 was preincubated overnight at 4°C with competition peptide (branched REIRR peptide) and negative control peptide (branched TGGVK peptide) at the indicated ratio of molar concentration.

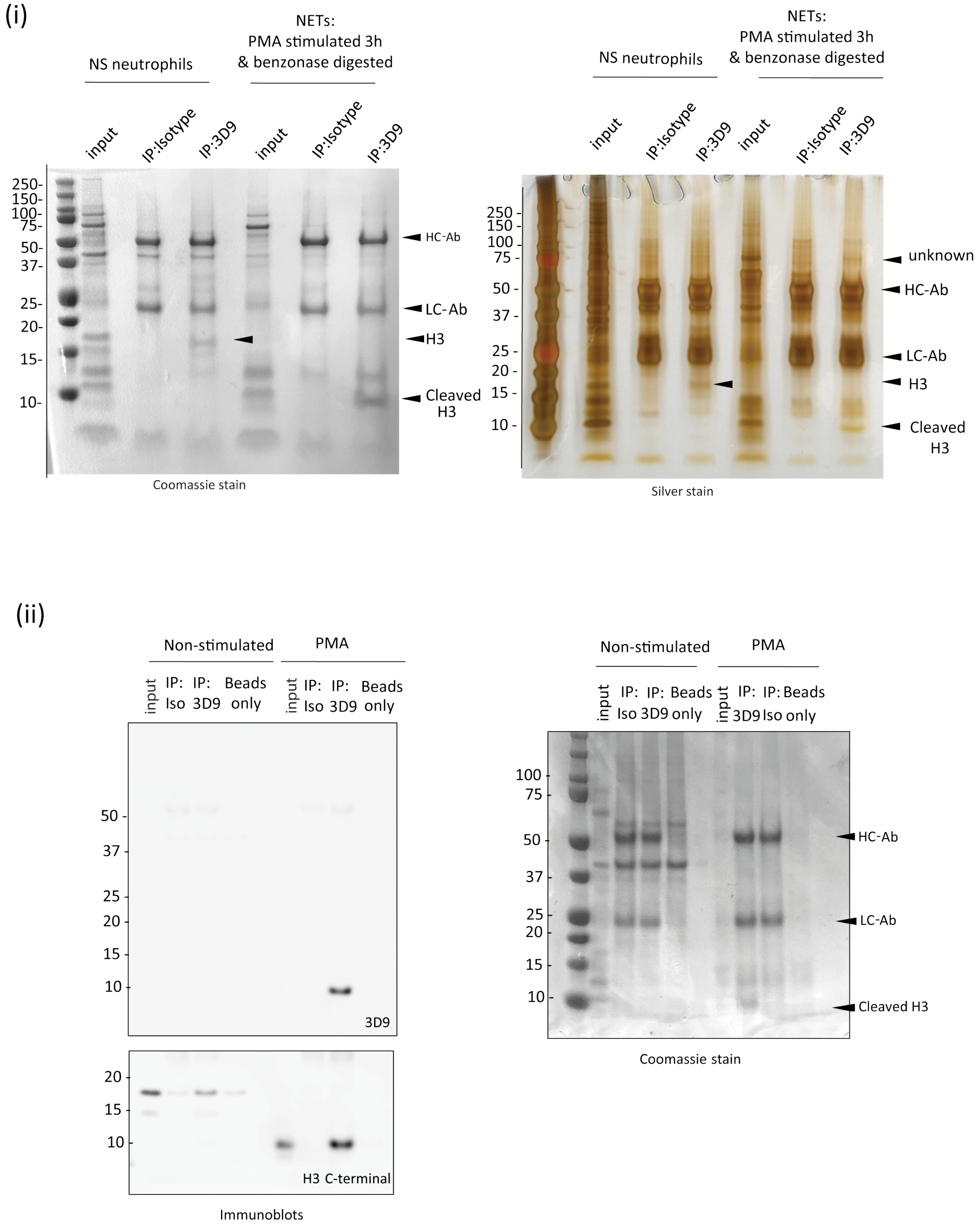


**Figure 3-figure supplement 4. 3D9 immunoprecipitation**

Neutrophils were seeded in 10 ml petri dishes and stimulated for 3h as indicated. Before total cell lysis and immunoprecipitation, NETs were digested with benzonase to liberate histones and octamers for antibody binding. (**i**) Coomassie stained gel of proteins immunoprecipitated by 3D9 or isotype control and the corresponding silver stained gel. The predicted cleaved H3 and intact H3 bands are indicated. HC-Ab: heavy chain of antibody, LC-Ab: light chain of antibody. (**ii**) Immunoblot for total H3 (C-terminal antibody) and cleaved H3 of the proteins immunoprecipitated by 3D9 or isotype control antibodies and Coomassie stained gel of samples run in parallel. Gels and blots are representative of 3 independent experiments.

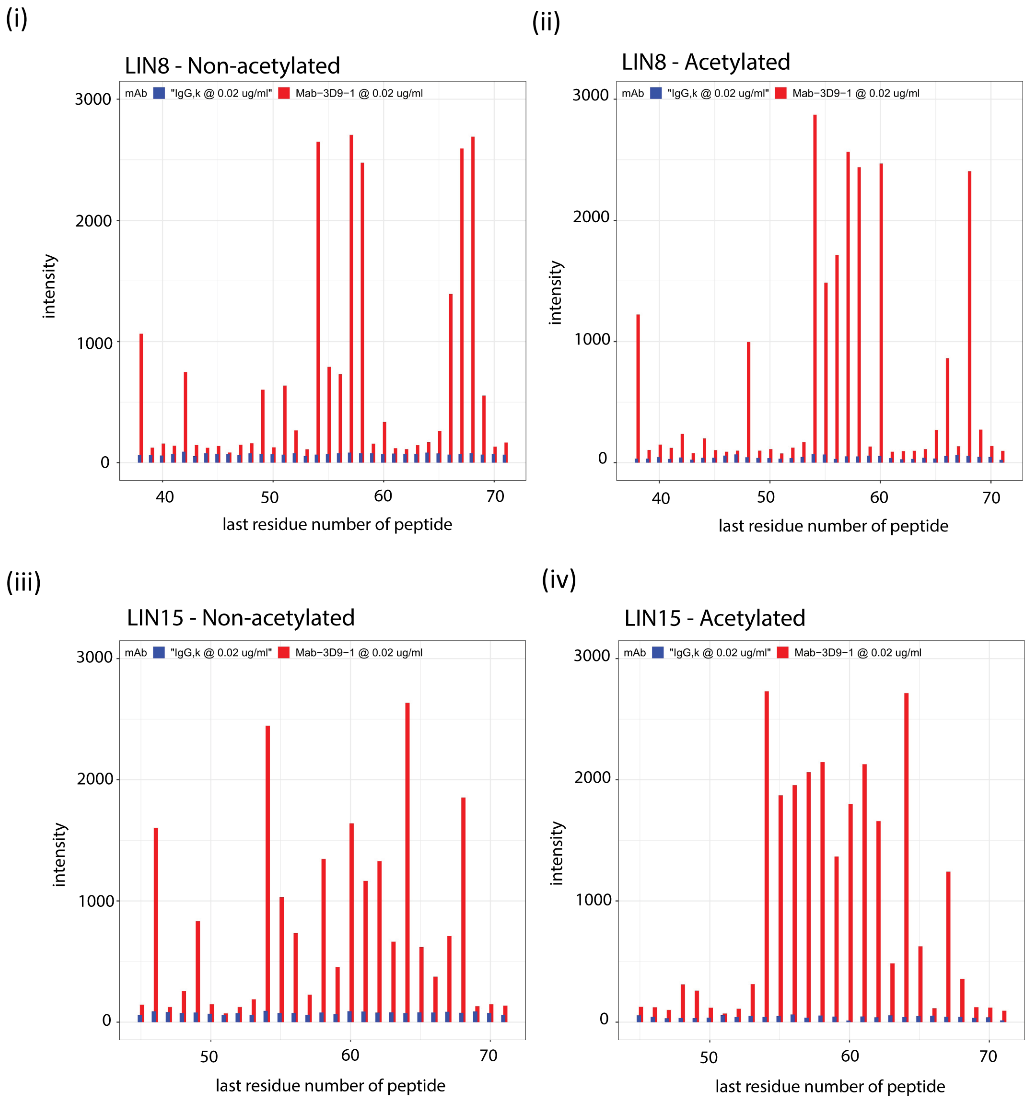


**Figure 4-figure supplement 1. Binding profiles recorded for 3D9 on the linear peptide array.** Antibodies 3D9 (red) and isotype control IgG,k (blue) were incubated on arrays with overlapping linear 8-mer (LIN8 – [**i, ii**]) and 15-mer (LIN15 – [**iii, iv**]) peptides and binding measured by ELISA. Separate arrays were used with peptides that were acetylated (**ii, iv**) or non-acetylated (**i, iii**) at the N-terminus. Signal intensities are plotted on the y axis and positions of the last residues of a peptide with respect to the target sequence is on the x axis. The antibodies were tested with the arrays 3 or more times. Peptides are listed in Supplementary file 1. Source data can be found in Figure 4-figure supplement 1-Source Data 1.

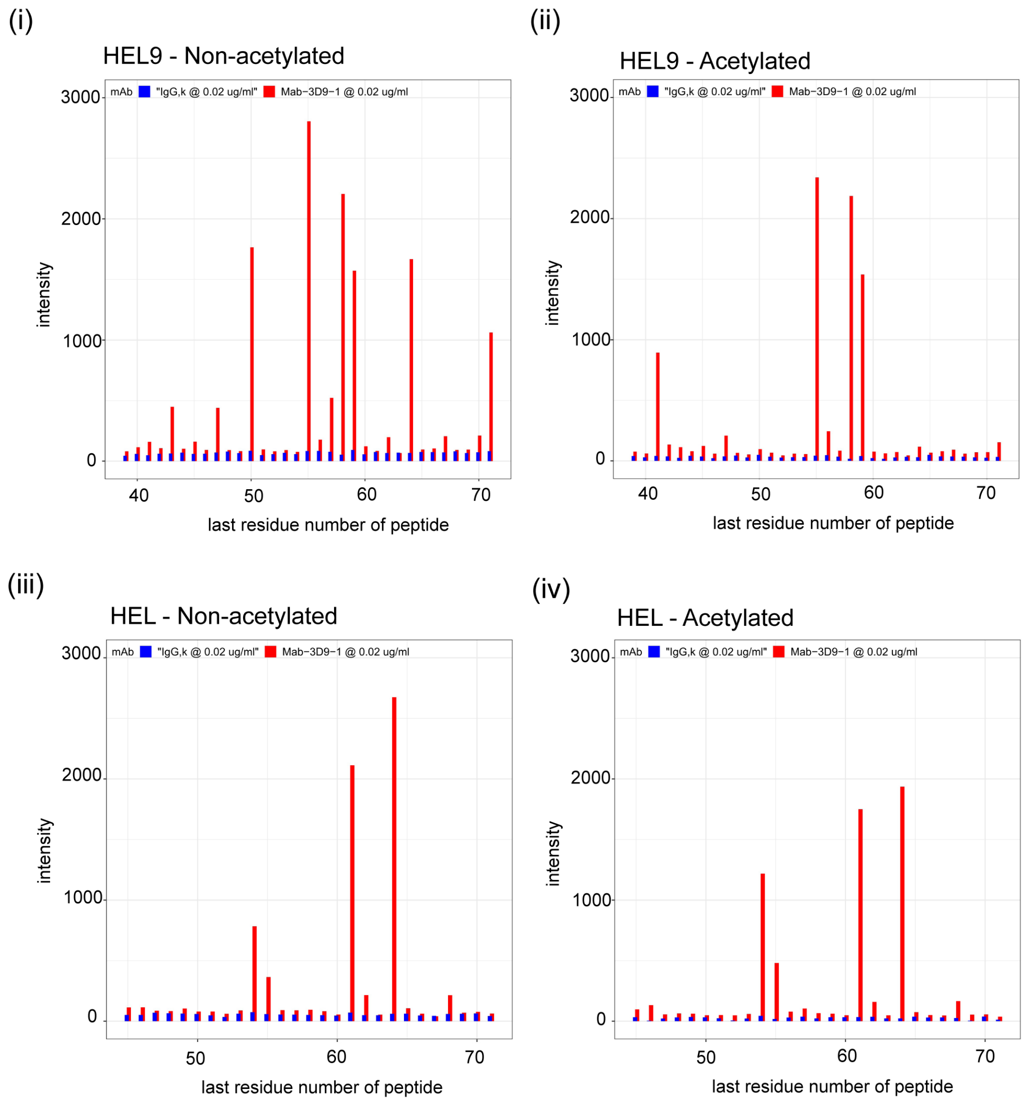


**Figure 4-figure supplement 2. Binding profiles recorded for 3D9 on helical peptide mimics arrays.** Antibodies 3D9 (red) and isotype control IgG,k (blue) were incubated on arrays with overlapping helical 9-mer (HEL9; [**i, ii**]) and 15-mer (HEL; [**iii, iv**]) peptides. Separate arrays were used with peptides that were acetylated (**ii, iv**) or non-acetylated (**i, iii**) at the N-terminus. Signal intensities are plotted on the y axis and positions of the last residues of a peptide with respect to the target sequence is on the x axis. The antibodies were tested with the arrays 3 times or more times. Peptides are listed in Supplementary file 1. Source data can be found in Figure 4-figure supplement 2-Source Data 1.

**S. Table 3: Summary of identified 3D9 binding regions in the peptide array**

| *Sample* | *Peptide type* | *Peak* | *Candidate epitope* |
|---|---|---|---|
| 3D9 | LIN8 | 47-58 | REIRRYQK |
| | | | VALREIRR |
| | | | EIRRYQKS |
| | | 60-68 | LLIRKLP |
| | | | ELLIRKLP |
| | LIN8 Ac | 47-60 | VALREIRR |
| | | | REIRRYQK |
| | | | RRYQKSTE |
| | | | EIRRYQKS |
| | | | LREIRRYQ |
| | | 61-68 | LLIRKLPF |
| | LIN15 | 40 -64 | HRYRPGTVALREIRR |
| | | | REIRRYQKSTELLIR |
| | | | TVALREIRRYQKSTE |
| | | 1-33 | RYQKSTELLIRKLPF |
| | LIN15 Ac | 12-44 | HRYRPGTVALREIRR |
| | | | REIRRYQKSTELLIR |
| | | | PGTVALREIRRYQKS |
| | | | VALREIRRYQKSTEL |
| | | | RPGTVALREIRRYQK |
| | | | YRPGTVALREIRRYQ |
| | | | RYRPGTVALREIRRY |
| | | | TVALREIRRYQKSTE |
| | | | ALREIRRYQKSTELL |

**Figure 4-figure supplement 3. S.Table 3. Summary of identified 3D9 binding regions in the peptide array.**
Epitope candidates are listed per peak (indicated by residue numbers) and are in order of intensity. The highest binder within a peak is at least in the 3rd quartile for the peptide type. Putative core epitopes are indicated in grey. Core epitopes were determined based on common sequences in the overlapping peptide sequences (peaks within 40% of the top peak).

(i)

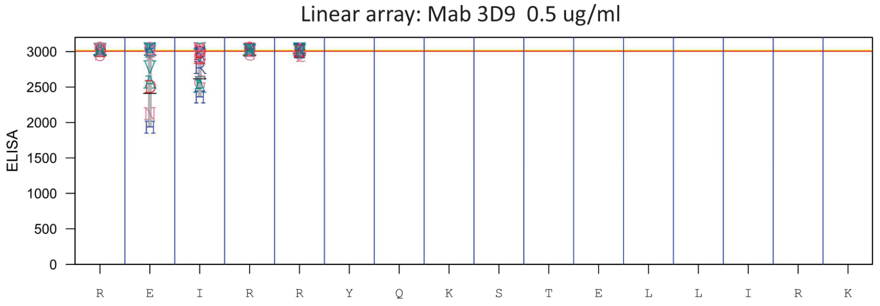


(ii)

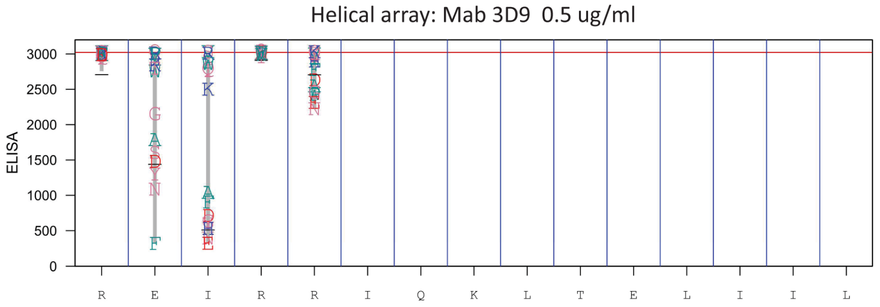


**Figure 4-figure supplement 4. Fine epitope mapping by replacement analysis.**
Linear (**i**) peptides were generated bearing single amino acid substitutions at each position of the native peptide sequence REIRRYQKSTELLIRK of histone H3. In addition, helical peptide mimics were made for the replacement analysis (**ii**). The peptides were left unacetylated to mimic the cleaved free N-terminus of the peptide. Antibody binding was analyzed at 0.5 µg/ml in PBST + 10% SQ (proprietary buffer). Values obtained for replacements are indicated by the letter code for each replacement residue plotted at the height of the recorded value at a given position. At the bottom of the plot the native sequence is indicated. Red line indicates median value of the native peptide sequence data. Source data can be found in Figure 4-figure supplement 4-Source Data 1.

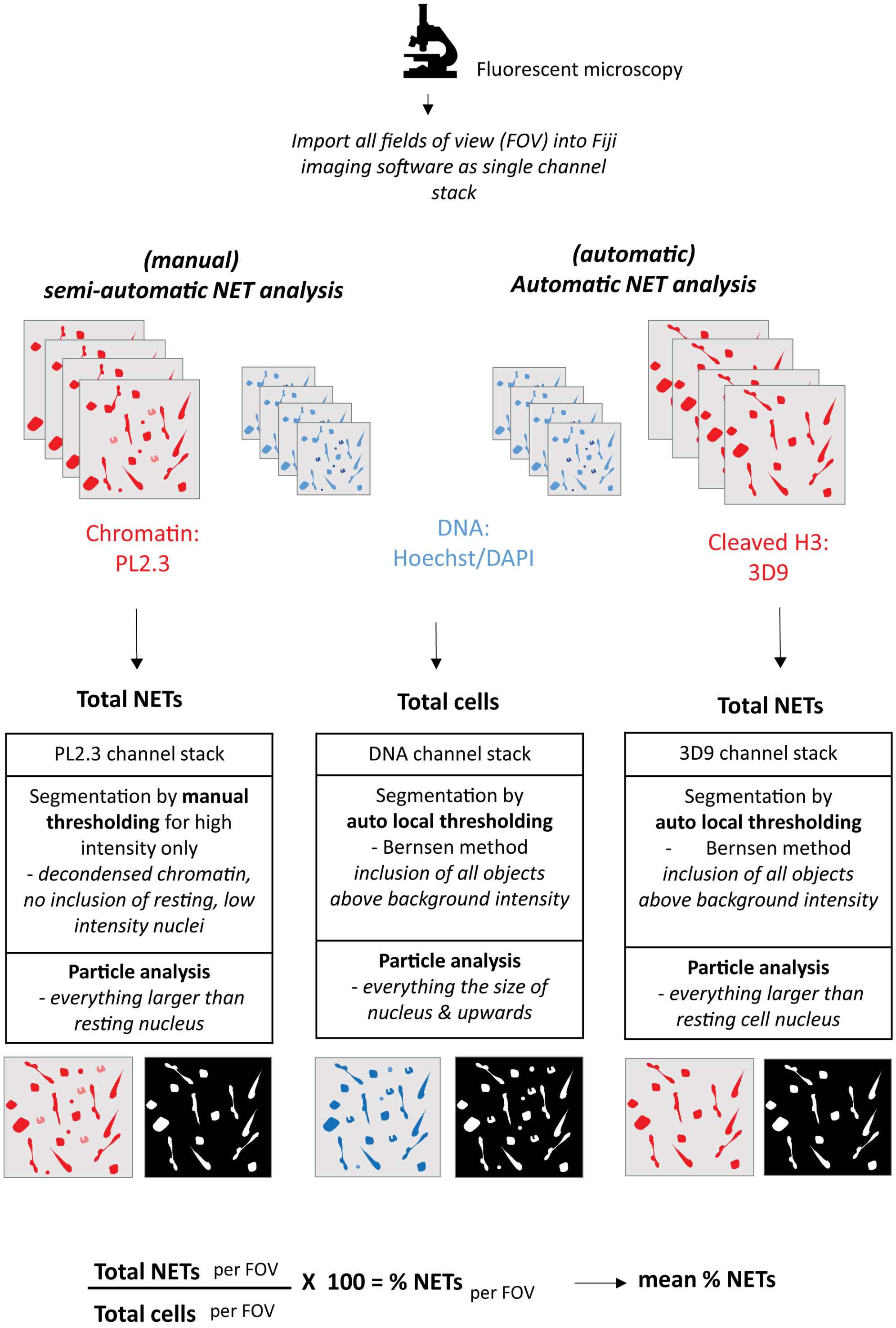


**Figure 5-figure supplement 1. Workflow of NET analysis methods**
Schematic of imaging, segmentation, thresholding and particle analysis for both automatic and semi-automatic NET quantification.

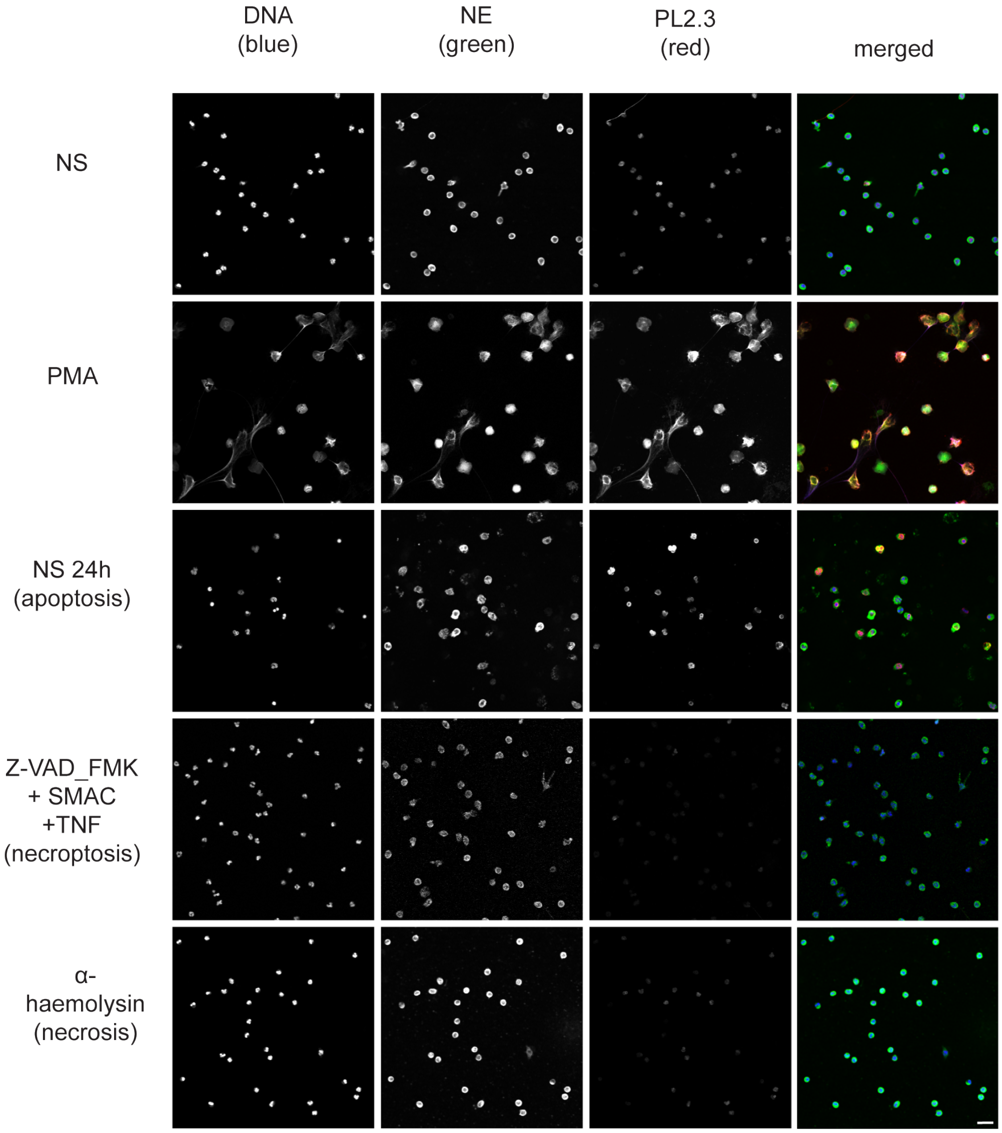


**Figure 8-figure supplement 1. Comparison of PL2.3 detection in response to apoptotic, necroptotic & necrotic cell death stimuli.**
Confocal immunofluorescent microscopy of neutrophils stimulated with different cell death stimuli and subsequently stained with Hoechst, anti-neutrophil elastase (NE) and PL2.3. NETs were induced with PMA (100 nM, 3h). Apoptosis was induced in resting neutrophils by incubation for 24h without stimulation. Neutrophils were stimulated with Z-VAD-FMK (50 µM) plus SMAC mimetic (100 nM) plus TNF (50 ng/ml) for 6h to induce necroptosis. Necrosis was induced with the pore forming toxin α-haemolysin (25 µg/ml). Images were taken at 20X and are representative of 3 independent experiments. Scalebar 20 µm. A comparison was made with parallel samples stained with the cleaved histone antibody 3D9 and are presented in Figure.8.

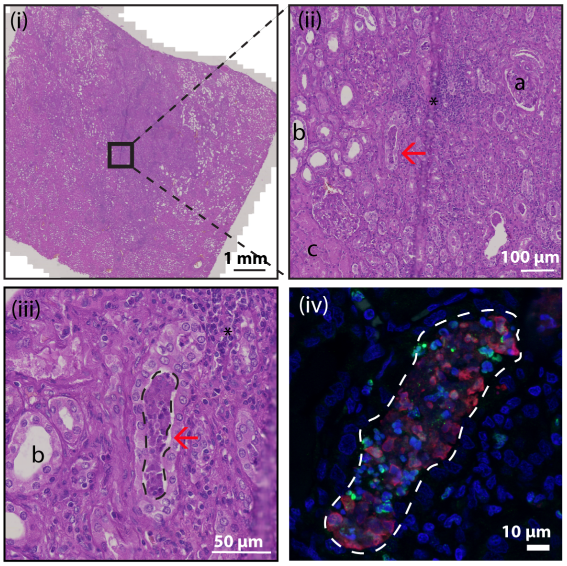


**Figure 9-figure supplement 1. Hematoxylin & eosin (HE) stain of kidney section**
HE stained tissue overview of NETs represented in Figure.9B and C. Black box indicates area for further magnification in images (ii) and (iii). The red arrow indicates the location of the NET in the collecting duct. 'a' - glomerulus, 'b'- tubule, 'c' - area of necrotic tubules and '*' – area of infiltrating lymphocytes (inflammation). Dashed line in (iii) and (iv) indicates area of 3D9 (red) and H3cit (green) staining from Fig 9C.

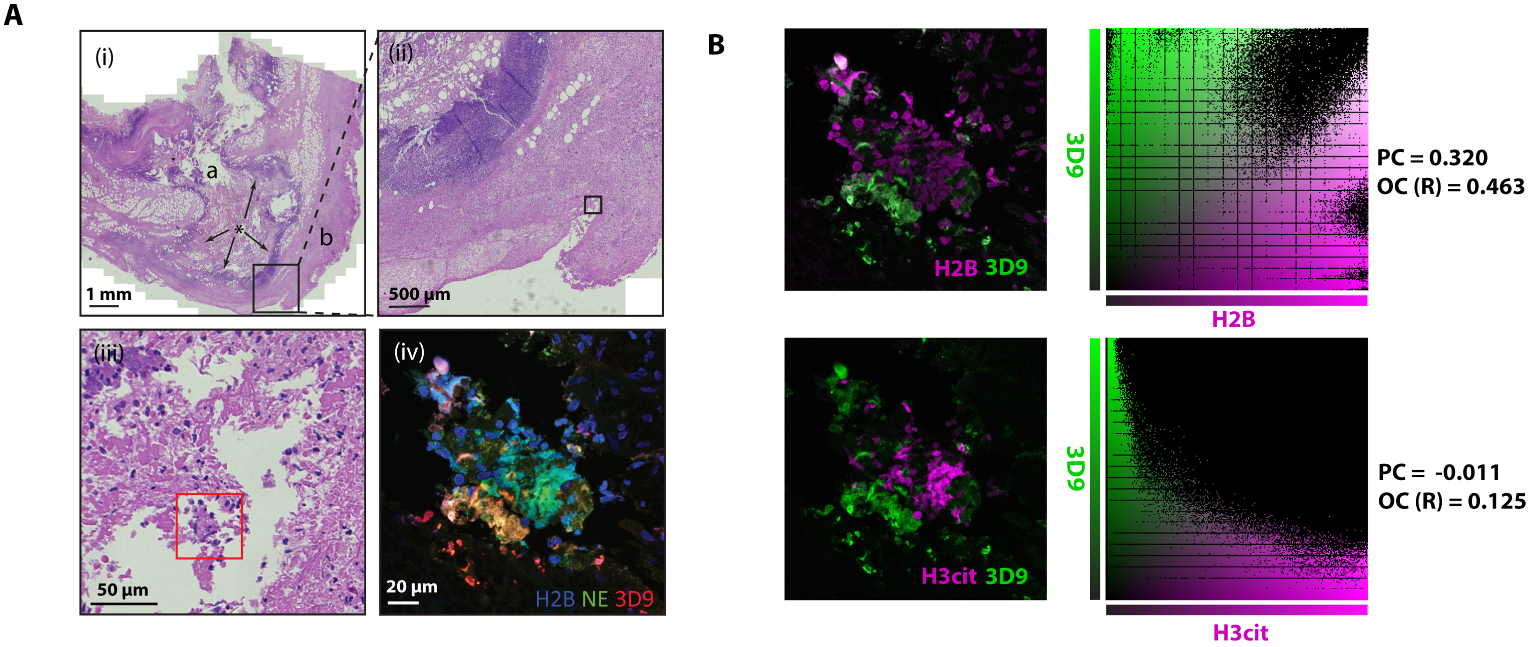


**Figure 10-figure supplement 1. Hematoxylin & eosin (HE) stain of gallbladder & colocalization analysis**
(**A**) HE stained overview of gallbladder stained for NETs in Figure 10. Magnification increases from (i)-(iii) and boxes represent the field of view of the next image. Red box indicates the area imaged by fluorescent confocal microscopy in Fig 9 (iv). 'a' – lumen; 'b' – serosa (exterior); '*' - area of infiltrating lymphocytes (inflammation). **(B)** Colocalisation analysis of 3D9 staining versus H2B staining and 3D9 versus H3cit. Analysis was performed using Volocity software and both the image analysed and associated scatter plot of pixel intensities are presented. PC = Pearsons Coefficient and OC (R)= overlap coefficient (R).

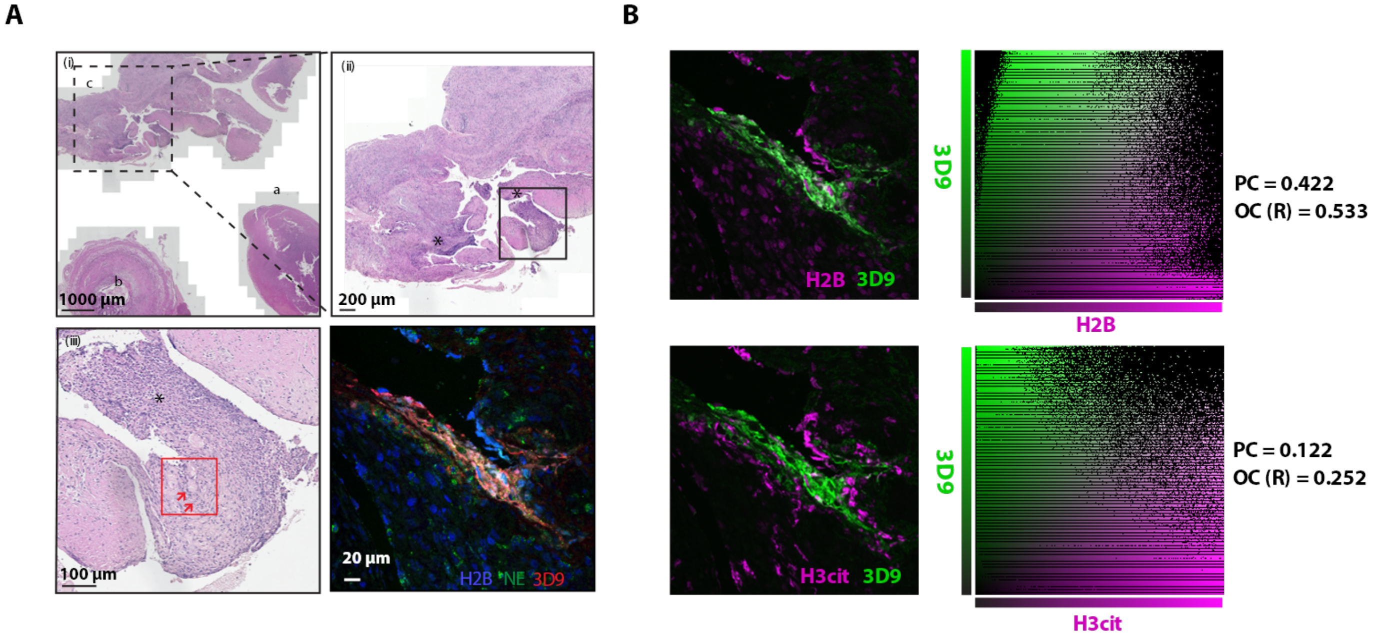


**Figure 11-figure supplement 1. Hematoxylin & eosin stain of inflamed appendix & colocalization analysis.**

(**A**) HE stained overview of appendix with appendicitis and peri-appendicitis stained for NETs in Fig 10. Magnification increases from (i)-(iii) and boxes represent the field of view of the next image. Red box indicates the area imaged by fluorescent confocal microscopy in Figure 11. NETs are indicated by arrows. a- appendix tip; b – lumen; c - associated inflamed tissue with peri-appendicitis. (**B**) Colocalisation analysis of 3D9 staining versus H2B staining and 3D9 versus H3cit. Analysis was performed using Volocity software and both the image analysed and associated scatter plot of pixel intensities are presented. PC = Pearsons Coefficient and OC (R) = overlap coefficient (R).
