## Supplemental File 2_methods for "Histone H3 clipping is a novel signature of human neutrophil extracellular traps"

### Supplemental methods

#### *Cytotoxicity assay*

To assess any toxicity of inhibitors LDH release was assessed using the CytoTox 96® Non-Radioactive Cytotoxicity Assay (Promega), according to the manufacturer's instructions. Neutrophils were seeded in RPMI in a 6 well plate with $1x10^5$ cells/100 µl. Inhibitors were preincubated with cells for 30 min at 37 °C at concentrations indicated in the figures. Assay was performed in triplicate.

#### *Reactive oxygen species assay*

Reactive oxygen species (ROS) production was measured using a luminol-HRP assay as described by Amulic *et al* (2017). Additionally, as all stages of the assay were performed in atmospheric conditions, normal tissue culture media was replaced with a carbonate and phenol red-free RPMI supplemented as before (Seahorse XF RPMI Medium #103336-100, Agilent). Inhibitors were preincubated with cells for 30 min at 37 °C at concentrations indicated in the figures. Immediately before stimulation, luminol (50 µM) and HRP (1.2 U/ml) were added followed by 50 nM PMA. Luminesce measurements were taken every 30 s using a luminescence plate reader (VICTOR Light luminescence counter, Perkin Elmer) and expressed as Relative Light Units (RLU). Assay was performed in triplicate.

#### *Mass spectrometry*

Protein spots were excised and transferred to 0.5 ml Eppendorf tubes. Samples were destained in 500 µl of 200 mM ammoniumbicarbonate (ABC) in 50% acetonitrile (ACN) for 30 min at 37°C and equilibration in 200 µl of 50 mM ABC, 5% ACN for 30 min at 37°C. The samples were then dried at room temperature for 60 min. Protein digestion was performed overnight at 37 °C with 100 ng trypsin (spots 1, 3 ,7 with 200 ng) in 25 µl of 50 mM ABC, 5% ACN. The supernatants were transferred into new Eppendorf tubes and additional peptides were extracted by applying 25 µl of 60% ACN, 0.5% trifluoroacetic acid (TFA) for 10 min, followed by 25 µl of 100% ACN for 10 min. All supernatants were combined and dried in an Eppendorf Concentrator at 45 °C.

After solubilization in 15 µl of 0.1% TFA, the peptides were desalted and concentrated with ZipTips and eluted with 1 µl alpha-cyano-4-hydroxycinnamic acid (5 mg/ml in 60% ACN, 0.3% TFA) onto the MALDI plate. Peptide mass fingerprints (PMF) and fragment spectra (MSMS) of the five most intense peaks were measured using a 4700 Proteomics Analyzer (AB Sciex). The database search was performed applying the Mascot MS/MS Ion Search function for searching the Swiss-Prot subset of human proteins. A peptide mass tolerance of 30 ppm and ± 0.3 Da for the fragment mass tolerance was allowed. One missed cleavage, oxidation of methionine, N-terminal acetylation of the protein, propionamide at cysteine residues and N-terminal pyroglutamic acid formation were defined as variable modifications. The following identification criteria were used: minimum 30% sequence coverage; or minimum 15% sequence coverage and one MS/MS confirmation; or sequence coverage below 15% and at least two MS/MS confirmations.

##### *Immunoprecipitation (IP) of clipped histones*

Magnetic Protein G coupled beads (Invitrogen) were washed (x3) with equal volume PBS. For each IP sample 20 µl of beads were prepared. As 3D9 was a mouse IgG1, a bridging antibody (anti-mouse, Active motif #53017) was used to improve binding to protein G couple beads. Bridging antibody (50 µg) was incubated with 20 µl of magnetic beads for 1 h at 4°C with gentle inversion. Beads were washed twice with PBS. Beads were then incubated with 50 µg 3D9 or isotype control (IgG1, k - Genscript clone 1H7A8) for 3h at RT (with gentle rotation). Beads were washed x3 and resuspended in 2 ml of PBS in a 15 ml falcon tube. To permanently crosslink the antibodies, PFA fixation was used. 2ml of 1% PFA (in PBS) was added to the beads and vortexed immediately for 1 min. The reaction was then quenched with 125 mM glycine (pH 8.0) and incubated on ice for 5 min. Beads were then washed x5 with IP lysis buffer (20 mM Tris, 150 mM NaCl, 1% Triton X-100, pH 7.5) and the blocked for 1 h at RT with 1% BSA in IP lysis buffer. Beads were used immediately or stored overnight at 4°C. The next day, neutrophils (1 x$10^7$) were seeded in Petri dishes with 10 ml of medium, left unstimulated or stimulated with

100 nM PMA for 3h at 37 °C. After stimulation, cells/NETs were washed (x3) very gently with equal volume of PBS (plus 1mM AEBSF). After the last wash NETs were digested in 1 ml of Benzonase buffer (20 mM Tris-HCl, pH 7.6, 2 mM $MgCl_2$, 1 mM $CaCl_2$) with 250U benzonase (Sigma E1014) and 1 mM AEBSF, 20 µM NEi and CGi for 30 min at 37°C, with intermittent rocking to distribute the buffer. The reaction was stopped with the addition of 4 mM EDTA and 4 mM EGTA. Cells were further lysed with the addition of 300 µl of 5X IP lysis buffer (plus 5X protease inhibitors). Dishes were incubated on ice for 10 min. Cells and NETs were then collected by scraping into an Eppendorf tube. For multiple dishes at the same time point and stimulation, the samples were pooled and divided for the IP step which was performed immediately. For each IP, lysate from $1x10^7$ cells was used with 15 µl of coupled beads and incubated overnight at 4°C with gentle rotation. The next day beads were washed 5 times with IP lysis buffer plus inhibitors. After the final wash beads were resuspended in 30 µl 1X SDS-PAGE sample loading buffer. Samples were analysed by SDS-PAGE and stained or immunoblotted as indicated. Coomassie Instant blue stain or Thermofisher silver staining kit were used according to the manufacturer's instructions.

*Peptide competition assay*

Neutrophils were seeded on coverslips in a 24 well dish ($1x10^5$ cells/well) and stimulated with PMA for 3h and fixed with 2% PFA. Prior to staining, 3D9 (2 ug/ml = ~12.9 nM) was preincubated overnight with rotation at 4°C in PBS with 0.01-200X fold molar excess of branched REIRR peptide. TGGVK branched peptide was used as negative control. The next day competed antibody-peptide solutions were diluted 1:1 with blocking buffer and the immunostaining procedure was followed as before. Peptide competition was assessed visually by fluorescent microscopy.

*Epitope mapping*

Antibody binding to human H3 was assessed by a combination of linear, conformational and amino acid replacement analysis epitope mapping with peptide synthesis and ELISA assays performed by Pepscan Presto B.V. (Leiden, The Netherlands). To reconstruct epitopes of the target molecule (H3 residues 30-70 - PATGGVKKPHRYRPGTVALREIRRYQKSTELLIRKLPFQRL) a library of peptide-based peptide mimics (Supplemental file 1) was synthesized using Fmoc-based solid-phase peptide synthesis. An amino functionalized polypropylene support was obtained by grafting with a proprietary hydrophilic polymer formulation, followed by reaction with t-butyloxycarbonyl-hexamethylenediamine (BocHMDA) using dicyclohexylcarbodiimide (DCC) with N-hydroxybenzotriazole (HOBt) and subsequent cleavage of the Boc-groups using trifluoroacetic acid (TFA). Standard Fmoc-peptide synthesis was used to synthesize peptides on the amino-functionalized solid support by custom modified JANUS liquid handling stations (Perkin Elmer). Synthesis of structural mimics (conformational analysis) was done using Pepscan's proprietary Chemically Linked Peptides on Scaffolds (CLIPS) technology. The binding of antibody to each of the synthesized peptides was tested in a Pepscan-based ELISA. The peptide arrays were incubated with primary antibody (3D9 or isotype control) at 0.02 µg/ml for linear and conformational analysis and 0.5 ug/ml for amino acid replacement analysis (overnight at 4°C). After washing, the peptide arrays were incubated with a 1/1000 dilution of rabbit anti-mouse IgG(H+L) HRP conjugate (Southern Biotech, Birmingham, AL, USA) for one hour at 25°C. After washing, the peroxidase substrate 2,2'-azino-di-3- ethylbenzthiazoline sulfonate (ABTS) and 20 µl/ml of 3 % H2O2 were added. After one hour, the colour development was quantified using a charge coupled device (CCD) - camera and an image processing system. The values obtained from the CCD camera ranged from 0 to 3000 mAU, similar to a standard 96-well plate ELISA-reader. The results were quantified and stored in the Peplab database. To verify the quality of the synthesized peptides, a separate set of positive and negative control peptides was synthesized in parallel. These were screened with commercial antibodies 3C9 and 57 (Posthumus et al., 1990).

To evaluate the binding and determine the core epitope, screenings were optimized for concentration and blocking conditions for each sample until clear signals over background values were observed. Peaks were defined as at least in the third quartile for the specific peptide mimic values. In addition, a binding event was only noted if multiple overlapping peaks were present within the binding region. To determine a core epitope, peptides within 40% of the main peak intensity were evaluated.
